# Architecture and infection-sensing mechanism of the bacterial PARIS defense system

**DOI:** 10.1101/2024.01.02.573835

**Authors:** Amar Deep, Qishan Liang, Eray Enustun, Joe Pogliano, Kevin D. Corbett

## Abstract

Bacteria and the viruses that infect them (bacteriophages or phages) are engaged in an evolutionary arms race that has resulted in the development of hundreds of bacterial defense systems and myriad phage-encoded counterdefenses^1–5^. While the mechanisms of many bacterial defense systems are known^1^, how these systems avoid toxicity outside infection yet activate quickly upon sensing phage infection is less well understood. Here, we show that the bacterial <u>P</u>hage <u>A</u>nti-<u>R</u>estriction-<u>I</u>nduced <u>S</u>ystem (PARIS) operates as a toxin-antitoxin system, in which the antitoxin AriA sequesters and inactivates the toxin AriB until triggered by the T7 phage counterdefense protein Ocr. Using cryoelectron microscopy (cryoEM), we show that AriA is structurally similar to dimeric SMC-family ATPases but assembles into a distinctive homohexameric complex through two distinct oligomerization interfaces. In the absence of infection, the AriA hexamer binds up to three monomers of AriB, maintaining them in an inactive state. Ocr binding to the AriA-AriB complex triggers rearrangement of the AriA hexamer, releasing AriB and allowing it to dimerize and activate. AriB is a toprim/OLD-family nuclease whose activation arrests cell growth and inhibits phage propagation by globally inhibiting protein translation. Collectively, our findings reveal the intricate molecular mechanisms of a bacterial defense system that evolved in response to a phage counterdefense protein, and highlight how an SMC-family ATPase has been adapted as a bacterial infection sensor.

## Introduction

The ongoing arms race between the bacteria and phage has driven the development of an extensive array of innate and adaptive immune strategies in bacteria^3,4^, as well as counter-immune strategies in phages^1,6^. In addition to the well-known restriction-modification (RM) and CRISPR-Cas systems, over one hundred other systems have been identified that target specific phage components or disrupt key metabolic pathways in the host cell to prevent phage propagation^3–5,7^. A typical bacterial genome harbors 5-20 such defense systems, largely grouped into “defense islands” in their genomes that show hallmarks of mobility through transposable elements and phage-mediated genetic transduction^1,8,9^. These systems likely provide overlapping or synergistic defense^10^, with first-responder systems targeting the incoming phage for immediate destruction without affecting the propagation of the host cell, and reserve systems that stall host cell growth through metabolite depletion, or kill the host cell to prevent phage propagation in a mechanism called abortive infection^11,12^.

Many phages encode factors that directly counter bacterial defense systems to enable infection of a host cell. In addition, phages can also carry defense systems of their own that prevent “superinfection” by other phages^5,13,14^. PARIS (Phage Anti-Restriction-Induced System) is a defense system that was initially identified in a P4-like phage satellite locus integrated into host *E. coli* genomes, and is also found in defense islands in bacterial genomes^5^. PARIS consists of a predicted ATPase protein (AriA) and a predicted toprim/OLD-family nuclease (AriB), and has been shown to act through abortive infection^5^. A striking feature of PARIS is that the system is activated by a T7 phage-encoded protein, Ocr, which has been shown to inhibit RM and Bacteriophage Exclusion (BREX) systems by mimicking the structure of DNA and binding to defense proteins^15–17^. Thus, PARIS can be considered a reserve defense system that is triggered specifically when first-responder defense systems are compromised by Ocr.

Here we reveal that PARIS functions as a type II toxin-antitoxin (TA) system with the AriA antitoxin binding and sequestering the toxin AriB in the absence of infection. In the presence of Ocr, AriA releases AriB through conformational changes coupled to Ocr binding and ATP hydrolysis, enabling the activation of AriB. Activated AriB induces cellular growth arrest by blocking protein translation, thereby inhibiting phage propagation. These data reveal the detailed structural mechanisms of a bacterial defense system triggered by a phage-encoded anti-restriction factor and provide insight into the broader family of toprim/OLD nuclease-containing defense systems in bacteria.

## Results

### PARIS is a type II toxin-antitoxin system

To understand the mechanistic basis of antiphage protection by PARIS, we first examined the *E. coli* B185 PARIS system, which comprises the AriA and AriB proteins (Figure 1a and Supplementary Figure 1a) and was previously shown to protect its host against diverse phages including T4 and T7^5^. While AriA was originally predicted to encode a AAA+ family ATPase^5^, our sequence analysis and Al-phaFold-based structure predictions (not shown) suggest that it is instead an ABC ATPase related to Rad50 and SMC-family proteins. Supporting the idea that AriA’s predicted ATPase activity is important for the phage defense activity of PARIS, we found that an AriA Walker B motif mutant (E393Q; AriA^EQ^), which eliminates ATP hydrolysis by AriA (Supplementary Figure 1b-c), also eliminates protection against phage T7 (Figure 1b and Supplementary Figure 1d). AriB is predicted to encode a metal-binding domain termed toprim (<u>top</u>oisomerase-<u>prim</u>ase) that is also found in OLD-family nucleases^5^, and we found that mutating a conserved residue in its predicted metal-binding site (E90A; AriB^E90A^) also eliminates protection against phage T7 (Figure 1b and Supplementary Figure 1d).

**Figure 1.**
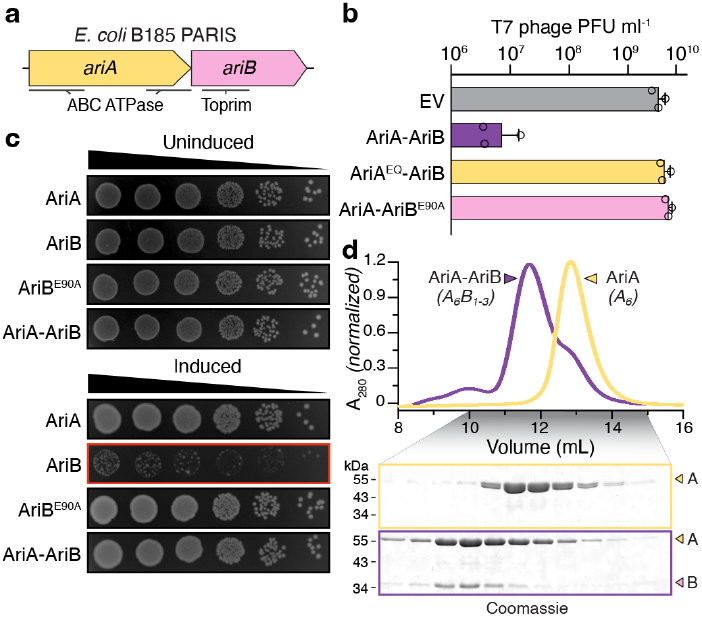
PARIS is a toxin-antitoxin system. **(a)** Schematic representation of the *E. coli* B185 PARIS system with predicted domain annotations for AriA and AriB proteins. **(b)** Analysis of EV (empty vector), AriA-AriB system (cloned as an operon), and their subunit mutants for their ability to defend against T7 phage infection. AriA^EQ^: E393Q Walker B motif ATPase-deficient mutant (see **Supplementary Figure 1c**). Data represent the mean plaque-forming units (PFU) mL^−1^ of phage T7 from three independent replicates, with individual data points shown (n = 3; see representative plaque assay in **Supplementary Figure 1d**). **(c)** Bacterial dilution spotting (10-fold) assay performed to measure the toxicity of arabinose-inducible AriA, AriB, AriA-AriB, and AriB^E90A^. A red outline indicates a condition where growth inhibition is observed. *Uninduced:* LB media + 0.4% glucose (suppressor) + antibiotics; *Induced:* LB media + antibiotics + 0.2% arabinose. See **Supplementary Figure 1e-g** for related bacterial growth curves. **(d)** Size exclusion chromatography analysis of purified AriA and AriA-AriB complex. The noted oligomeric states were determined using size-exclusion chromatography coupled with multi-angle light scattering (SEC-MALS) (see **Supplementary Figure 2**).

Based on the conserved operonic architecture of PARIS and the predicted functions of both proteins, we hypothesized that PARIS may comprise a toxin-antitoxin (TA) system with AriB as toxin and AriA as antitoxin. To test this idea, we overexpressed AriA, AriB, or both in *E. coli*, and measured the impact on bacterial growth. AriB expression strongly inhibited cell growth, and this effect was alleviated by coexpression of AriA (Figure 1c and Supplementary Figure 1e-g). Overexpression of AriB^E90A^ had no effect on bacterial growth (Figure 1c), suggesting that the observed growth arrest is a result of AriB’s predicted nuclease activity.

Bacterial TA systems have been classified into eight types based on whether the antitoxin is a protein or a non-coding RNA, and on how the antitoxin neutralizes the toxin^18^. To classify PARIS, we tested whether AriA and AriB physically interact. To avoid the toxicity associated with AriB expression, we employed AriB^E90A^ for the physical interaction analysis. Using size-exclusion chromatography coupled to multi-angle light scattering (SEC-MALS), we found that AriA forms a stable homohexamer (AriA_6_) that can bind up to three AriB monomers (AriA_6_B_3_; Figure 1d and Supplementary Figure 2a-c). Notably, expression of AriA and AriB from a native-like operon resulted in an excess of AriA and only partial occupancy of AriB (AriA_6_B_1-2_) (Supplementary Figure 2b); generation of the saturated AriA_6_B_3_ complex required separate expression of additional AriB (Supplementary Figure 2c). When combined with our finding that AriB expression on its own is toxic to *E. coli* cells, these data suggest that PARIS is a type II TA system in which the antitoxin AriA is produced in excess and directly binds and neutralizes the toxin AriB. Finally, while coexpression of AriB^E90A^ with AriA resulted in a soluble AriA-AriB complex, we were unable to express and purify AriB^E90A^ on its own. These data suggest that in addition to serving as an antitoxin, AriA likely serves as a chaperone for AriB, maintaining its folding and solubility in cells.

### Structural basis of AriA hexamer assembly

ABC ATPases, including membrane transporters and SMC ATPases, typically form homo- or heterodimers^19^. To understand the molecular basis for homohexamer assembly of AriA, we determined a 3.6 Å resolution cryoEM structure of AriA^EQ^ (Supplementary Figure 3 and Supplementary Table 1). 2D class averages revealed a three-fold symmetric assembly with three short “legs” extending from a central hub (Figure 2a-c and Supplementary Figure 3). The final refined map and model includes four copies of AriA that represent two-thirds of the hexameric assembly, with poorly-resolved density representing the remaining two AriA subu-nits (Figure 2c-d).

**Figure 2.**
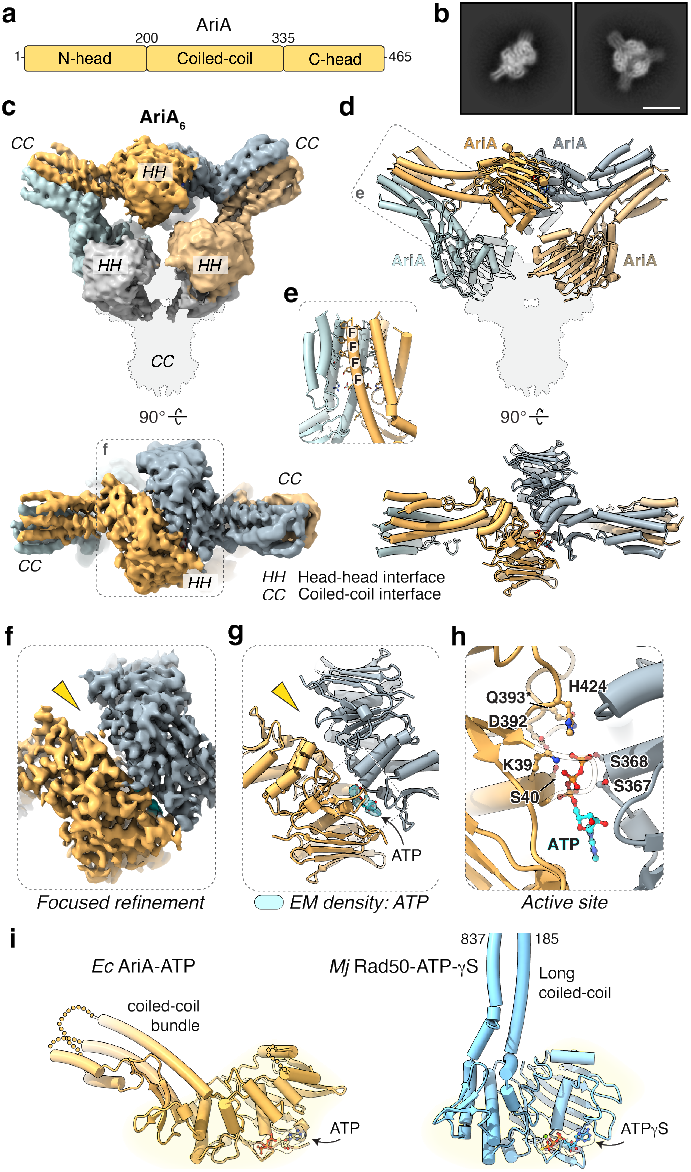
Structure of AriA. **(a)** Schematic representation of the domain structure of AriA. **(b)** Two 2D class averages from cryoEM analysis of the AriA^EQ^ sample, revealing a 3-fold pseudo-symmetric hexameric assembly (see **Supplementary Figure 3**). Scale bar = 10 nm. **(c)** Two views of the cryoEM density map of AriA^EQ^ illustrating a hexameric self-assembly, with four intact protomers in shades of yellow and blue, and the density of two poorly-resolved protomers in gray. CC and HH represent the coiled-coil and head-head dimer interfaces, respectively. **(d)** Cartoon representation of four ordered AriA protomers colored in shades of yellow and blue. **(e)** Inset from panel d, a cartoon view showing the CC interface, with residues involved in interaction depicted as stick representations for side chains. **(f)** A focused refinement map displaying the cryoEM density for the HH dimer. **(g)** Cartoon view of the AriA^EQ^ HH interaction interface, illustrating cryoEM density for a bound ATP molecule in cyan. The yellow arrow indicates an open site in the ATPase head region. **(h)** Close-up view of ATP bound to AriA. **(i)** Cartoon views of *E. coli* B185 AriA^EQ^ in yellow and *Methanocaldococcus jannaschii* Rad50 (PDB ID 3AV0), showing the region of structural alignment with a yellow highlight.

The AriA^EQ^ homohexamer is assembled through two distinct AriA-AriA interfaces we term “head-head” (HH) and “coiled-coil” (CC; Figure 2c-g). Each AriA protomer adopts a domain architecture similar to the ABC ATPase Rad50 and SMC-family ATPases, with N- and C-terminal ATPase subdomains separated by a coiled-coil region^20,21^. Unlike Rad50 and SMC ATPases whose coiled-coils dimerize at their distal ends through Zn^2+^-hook and hinge-hinge interactions, respectively, the coiled-coil region of each AriA protomer forms a short four-helix bundle and dimerizes along its length with a second AriA protomer through a hydrophobic interface involving four conserved phenylalanine residues (CC interface; Figure 2e). Compared to Rad50 and SMC ATPases, the coiled-coil of AriA also emerges from the ATPase domain at a dramatically different angle, enabling assembly of the AriA hexamer (Figure 2i and Supplementary Figure 4a-c).

The HH dimer interface of AriA resembles the canonical ATPase-domain dimer interface of Rad50 and SMC ATPases, in which two ATP molecules are typically sandwiched between the ATPase domains of opposite protomers^19^. In our structure of AriA^EQ^, the AriA^EQ^ ATPase-domain dimer is asymmetric, with one active site bound to ATP and the other active site open (Figure 2f-g). ATP binding is mediated through conserved residues in the Walker A and Walker B motifs of one protomer and the ABC ATPase signature motif from the opposite protomer (Figure 2h and Supplementary Figure 1b and 4d). ATPase assays confirmed that mutation of the AriA Walker A motif (K39I) and Walker B motif (D392A, Q393E) disrupted ATP hydrolysis (Supplementary Figure 1c). All mutant AriA proteins, including those designed specifically to disrupt nucleotide binding (K39I and D392A), retained the ability to form hexameric assemblies (Supplementary Figure 5a-c).

### AriA sequesters AriB in an inactive state

To understand how AriA and AriB interact with one another, we determined a cryoEM structure of the AriA^EQ^-AriB^E90A^ complex to 3.1 Å resolution. (Supplementary Figure 6 and Supplementary Table 1). 2D class averages revealed the familiar three-fold symmetric AriA hexamer plus three additional protrusions located between the AriA coiled-coil “legs” (Figure 3a-b and Supplementary Figure 6c). The final refined map revealed four well-ordered AriA protomers (two CC dimers linked by an HH interface) and one AriB protomer bound at the HH interface of the central pair of AriA protomers (Figure 3c). Focused refinement of AriB and the neighboring AriA protomers revealed that AriB binds directly between the AriA ATPase heads, interacting asymmetrically with both protomers through its N-terminal domain (Figure 3d-e). The AriB-AriA interface is anchored by hydrophobic residues in the first α-helix of AriB that dock within the cavity formed at the AriA HH interface, and this interaction is stabilized by multiple charged residues on both proteins participating in a network of hydrogen bonding and salt bridge interactions (Figure 3f). The asymmetric binding of AriB to the AriA HH interface structurally resembles the interaction of bacterial SMC ATPases MukB and JetC with the C-terminal WHD domain of their cognate kleisin subunits^20^ (Supplementary Figure 7a-b). While kleisin C-WHD binding boosts ATP hydrolysis in SMC ATPases^22^, we find that AriB binding reduces the ATPase activity of AriA ∼4-fold (Supplementary Figure 7c), potentially due to structural alterations imposed by AriB on the AriA HH interface. Compared to the AriA^EQ^ structure, the HH interface in the AriA^EQ^-AriB^E90A^ complex is relatively closed, with one ATP molecule sandwiched tightly between AriA ATPase domains, and a second ATP bound to the opposite active site, which remains partially open (Supplementary Figure 7d).

**Figure 3.**
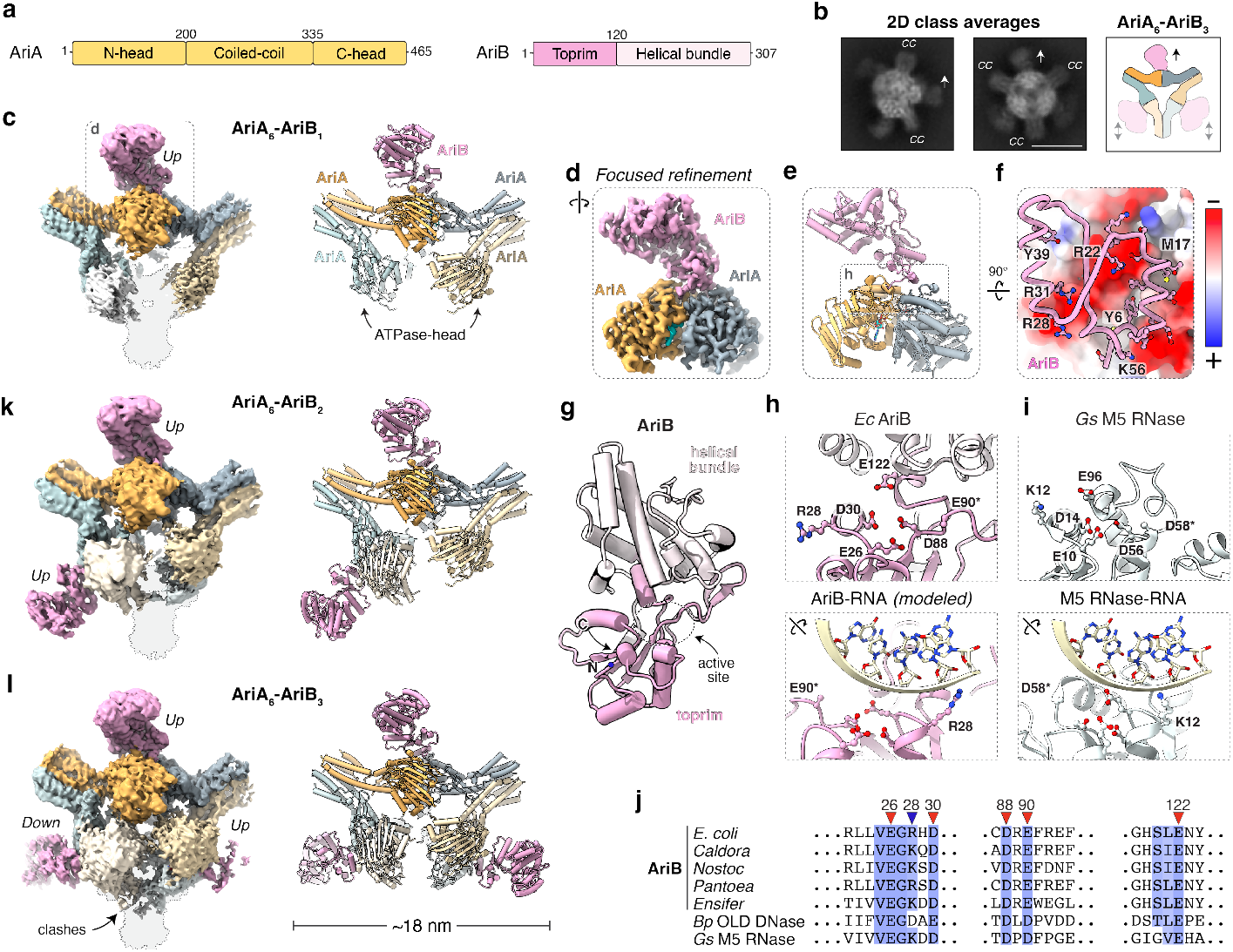
Structure of AriA-AriB complex. **(a)** Schematic representations of the domain structure of *E. coli* B185 AriA and AriB. **(b)** 2D class average from cryoEM analysis of AriA^EQ^-AriB^E90A^ along with a subunit schematic, revealing an AriA_6_B_3_ hetero-nonameric assembly. Scale bar = 10 nm. **(c)** cryoEM density map and cartoon representations of the AriA^EQ^-AriB^E90A^ complex. The density for the two missing AriA^EQ^ protomers is illustrated by a gray outline. AriA protomers are shown in shades of yellow and blue, while AriB is shown in light pink. **(d, e)** Focused refinement cryoEM map and model, colored as in panel c. **(f)** The inset shows AriB (cartoon) binding to the AriA HH interface (surface colored by electrostatic potential). **(g)** Cartoon representation of AriB. **(h)** *Top:* Close-up view of AriB active site, with conserved residues depicted in ball-and-stick representations for side chains. *Bottom:* A rotated close-up view of AriB’s active site with modeled RNA shown in gray. E90* denotes the AriB E90A mutation. **(i)** *Top:* Active site of *Geobacillus stearothermophilus* (*Gs*) M5 RNase (PDB ID 6TPQ), showing active site residues in ball-and-stick representations for side chains. *Bottom:* A rotated view of *Gs* M5 RNase (PDB ID 6TPQ) active site with RNA target shown in light-yellow^24^. See **Supplementary Figure 8a-d. (j)** Multiple sequence alignment close-up views of AriB and its structural homologs showing conservation of active site residues (*E. coli* AriB: NCBI accession # WP_000093097.1; *Caldora* sp. SIO3E6 AriB: NCBI accession # NES19199.1; *Nostoc* sp. PCC 7107 AriB accession # WP_015115038.1; *Pantoea* sp. YR343 AriB accession # WP_008101553.1; *Ensifer* unclassified AriB accession # WP_141687213.1; *Burkholderia pseudomallei* OLD DNase accession # WP_011852403.1; *Geobacillus stearothermophilus* M5 RNase accession # WP_095858694.1). **(k, l)** cryoEM density map and cartoon structural representations of the AriA_6_-AriB_2_ and AriA_6_-AriB_3_ complexes (colored as in panel c).

AriB comprises an N-terminal toprim-like nuclease domain and a C-terminal helical bundle domain (Figure 3g). The AriB toprim domain is structurally similar to the catalytic domains of OLD DNase from *Burkholderia pseudomallei* (*Bp*) and M5 RNase from *Geobacillus stearothermophilus* (*Gs*) (Figure 3h-i and Supplementary Figure 8a-d). Consistent with *Bp* OLD DNase^23^ and *Gs* M5 RNase^24^, AriB’s active site residues (E26, D30, D88, E90, and E122) are highly conserved and are positioned for metal ion coordination to cleave nucleic acids (Figure 3j and Supplementary Figure 8d). Comparative structural analysis suggests that AriB is more closely related to *Gs* M5 RNase, as both proteins lack a critical Lys residue found in *Bp* OLD (K562) that is required for DNA cleavage in this family of nucleases^23^ (Supplementary Figure 8d). Further, we identified another conserved positively charged residue, R28, whose equivalent in *Gs* M5 RNase (K12) interacts directly with the RNA backbone^24^ (Figure 3j and Supplementary Figure 8d). A charge-reversal mutant of this putative RNA binding residue (R28E) results in a loss of PARIS antiphage activity, suggesting that AriB’s inhibition of cell growth arises from the cleavage of as-yet unidentified nucleic acids (Supplementary Figure 8e).

Our SEC-MALS analysis showed that the AriA hexamer can bind up to three protomers of AriB. In our structural analysis, apart from the well-resolved central AriB protomer (Figure 3c), we detected weak density at the other two AriA HH interfaces that likely represent additional AriB protomers (Figure 3b). Considering how AriA and AriB interact, there are nine possible AriA-AriB assemblies that differ by AriB occupancy (one, two, or three AriA protomers bound) and orientation (arbitrarily assigned as “Up” or “Down”) on the AriA hexamer (Supplementary Figure 9). We generated structural templates for all nine possibilities and performed heterogeneous refinement in cryoSPARC with ∼500K particles, resulting in 3D reconstructions for each state at resolutions ranging from 3.1 Å to 6.2 Å (Figure 3k-l and Supplementary Figure 9). These structures show that PARIS can adopt both compositionally and conformationally heterogeneous forms, revealing dynamic variability and polymorphism within AriA-AriB complexes.

### The counterdefense protein Ocr triggers AriA-AriB dissociation

Prior work has shown that missense mutations in Ocr, a DNA-mimicking anti-restriction protein^15,16^, enable T7 phage to escape PARIS-mediated immunity^5^. We isolated mutant T7 phages that escape protection by PARIS, and found that three independent escaper clones all contain the same missense mutation in the gene encoding Ocr, mutating residue L82 to arginine (L82R; Supplementary Figure 10a-d). Both residues L82 and F54, which was previously shown to be mutated in PARIS escaper mutants^5^, lie at the dimer interface of Ocr, and we found by SEC-MALS that Ocr L82R is monomeric in solution (Supplementary Figure 10e). These data suggest that Ocr dimerization may be required to activate PARIS. We confirmed that coexpression of Ocr with AriA-AriB in *E. coli* cells causes growth arrest (Figure 4a and Supplementary Figure 11a-d). We also found that AriA-AriB+Ocr expression shows a consistently stronger growth arrest than expression of AriB alone (Supplementary Figure 11b-d), supporting the idea that AriA serves both as an antitoxin for AriB and as a chaperone to maintain AriB folding and solubility.

**Figure 4.**
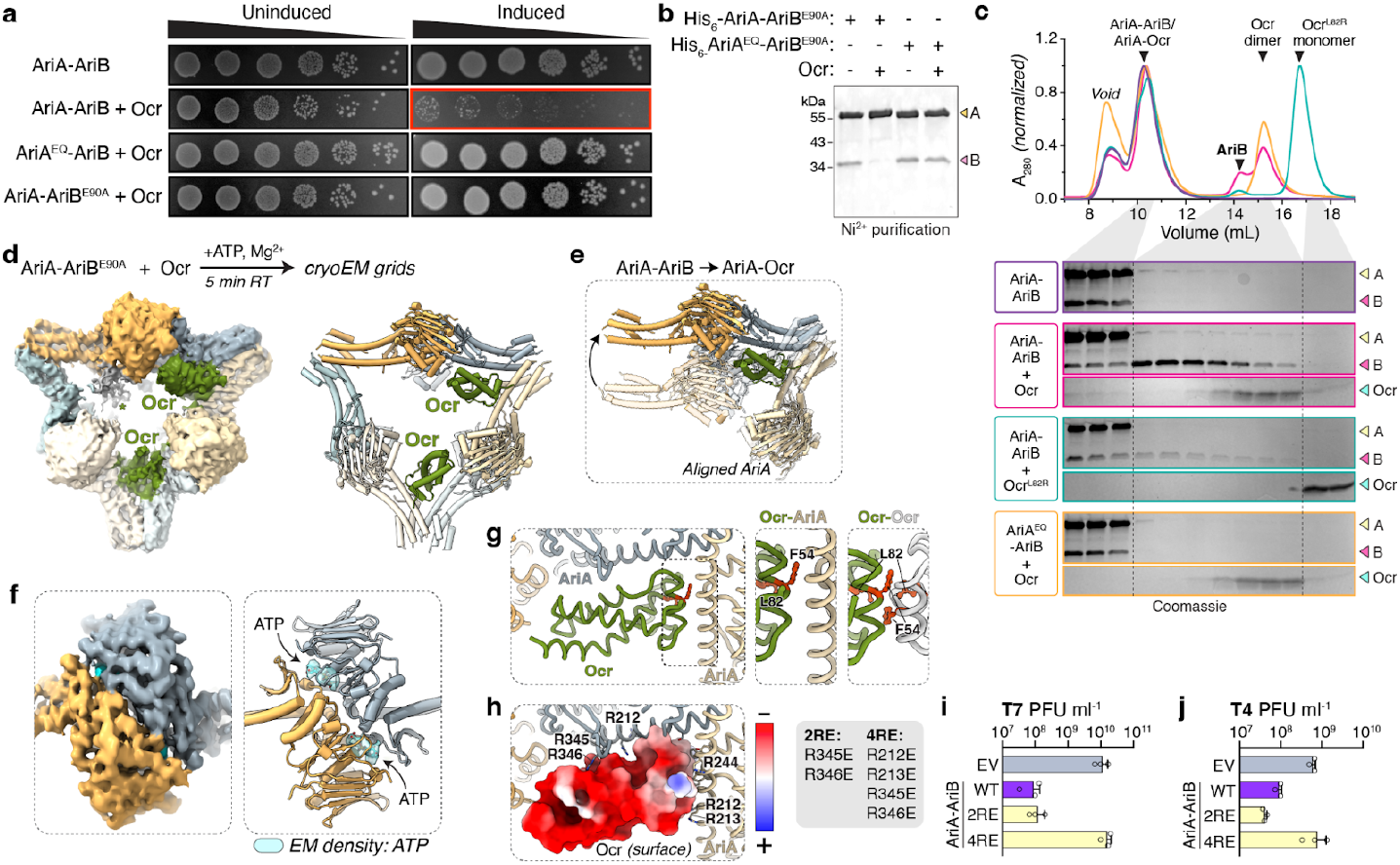
Structural basis of PARIS activation by T7 Ocr. **(a)** Bacterial dilution spotting (10-fold) assay performed to measure the toxicity of AriA-AriB, AriA^EQ^-AriB, and AriA-AriB^E90A^ upon co-expression with Ocr. *Uninduced:* LB media + 0.4% glucose (suppressor) + antibiotics; *Induced:* LB media + antibiotics + 0.2% arabinose (to induce AriA-AriB) + 0.1 mM IPTG (to induce Ocr). See **Supplementary Figure 11a** for induction with arabinose only, and **Supplementary Figure 11b-d** for related bacterial growth curves. **(b)** SDS-PAGE showing the results of coexpression of AriA-AriB with Ocr followed by Ni^2+^ affinity purification of AriA and associated proteins. **(c)** Size-exclusion chromatography analysis of AriA-AriB complexes upon mixing with Ocr. All mixtures were incubated in a buffer containing ATP and Mg^2+^ for five minutes at room temperature prior to column application). **(d)** *Top:* sample preparation strategy used to determine AriA-Ocr complex structure. *Bottom:* CryoEM density map and cartoon representations of the AriA-Ocr complex. AriA protomers are shown in shades of yellow and gray, consistent with coloring in previous figures. Two Ocr protomers are shown in green. An asterisk (*) represents EM density for a third Ocr protomer with partial occupancy. **(e)** Structural overlay showing conformational changes in AriA upon Ocr binding. AriA-AriB structure is shown in semi-transparent and AriA-Ocr is shown in solid color. For clarity, AriB and other AriA protomer chains are not shown. **(f)** CryoEM density map (left) and cartoon model (right) showing the AriA HH interface in the AriA-Ocr complex structure. The cryoEM density of bound ATP molecules is shown in cyan. **(g)** *Left:* Overall view of Ocr binding AriA, with subunits colored as in panel d. Right: closeup of the Ocr-AriA and Ocr-Ocr (PDB ID 1S7Z^16^) interfaces, showing the positions of Ocr residues F54 and L82 (colored red and shown as sticks). **(h)** View of the Ocr-AriA interface equivalent to panel h, with Ocr shown as an electrostatic surface and positively-charged AriA residues shown as sticks and labeled. **(i-j)** Analysis of EV (empty vector), Wt (AriA-AriB system, cloned as an operon), and AriA receptor pocket mutants for their ability to defend against T7 (panel i) and T4 (panel j) phage infections. 2RE and 4RE are the double and quadruple charge-reversal mutants of AriA in AriA-AriB operon (as shown in highlighted box on left). Data represent the mean plaque-forming units (PFU) mL^−1^ of phages T7 and T4 from three independent replicates, with individual data points shown (n = 3; see representative plaque assay in **Supplementary Figure 13c**).

To test the idea that Ocr activates PARIS by triggering dissociation of AriB from AriA, we coexpressed Ocr with His_6_ -AriA and AriB^E90A^ in *E. coli* and purified His_6_ -AriA and its associated protein(s) through Ni^2+^ affinity chromatography. We found that AriA was unable to bind AriB^E90A^ in the presence of Ocr, but that the AriA Walker B mutant E393Q was insensitive to Ocr and maintained AriB^E90A^ binding (Figure 4b). To assess whether Ocr can directly trigger AriA-AriB dissociation *in vitro*, we individually purified AriA-AriB^E90A^ and Ocr, mixed them in the presence of ATP, and performed size exclusion chromatography. We found that Ocr binds the AriA hexamer and at the same time triggers the release of AriB (Figure 4c). Consistent with our coexpression analysis, AriA^EQ^ was insensitive to Ocr, neither binding Ocr nor releasing AriB upon mixing (Figure 4c). Further, when we performed the same assay with the mutant Ocr^L82R^, we observed reduced but still detectable AriB release, but no stable Ocr^L82R^ binding to AriA (Figure 4c).

Together, these data suggest that Ocr binding to AriA triggers ATP hydrolysis and concurrent release of AriB. We found that at steady state, Ocr addition reduces AriA ATP hydrolysis by about half (Supplementary Figure 11e) and reduces the already-low ATP hydrolysis activity of the AriA-AriB complex by a similar amount (Supplementary Figure 11e). In contrast, Ocr^L82R^ had no effect on ATP hydrolysis by AriA but increased AriA-AriB ATP hydrolysis by about 50% (Supplementary Figure 11e). We interpret these data as indicating that both Ocr and AriB can inhibit AriA ATP hydrolysis at steady state, and that Ocr^L82R^ boosts steady-state ATP hydrolysis by triggering release of AriB, but then failing to stably associate with AriA.

### Structural basis of infection sensing by AriA

We next attempted to determine the structural basis for Ocr-triggered AriA-AriB dissociation. We combined AriA-AriB^E90A^ and Ocr in the presence of ATP, incubated for a short time, then froze samples for analysis by cryoEM (Supplementary Figure 12, Supplementary Figure 13a-b, and Supplementary Table 1). 2D class averages showed a three-fold symmetric assembly of AriA with no evidence of bound AriB (Supplementary Figure 12c). The final refined map showed a near three-fold symmetric assembly with six copies of AriA and two monomers of Ocr, with each Ocr monomer positioned between two AriA coiled-coil domains and wedging open the CC interface (Figure 4d). Ocr binding results in a significant outward motion of the AriA ATPase domains, and a consequent rearrangement of the HH interface (Figure 4e). In the AriA-Ocr structure, each HH interface is symmetric and bound to two molecules of ATP (Figure 4f).

While Ocr natively forms a homodimer (Supplementary Figure 10e), the protein binds AriA as a monomer. Ocr is positioned asymmetrically at the AriA CC interface, with the hydrophobic dimer interface of Ocr docking against an α-helix in one AriA coiled-coil domain. Strikingly, this interface structurally mimics the Ocr dimer interface (Figure 4g). Ocr residues F54 and L82, whose mutation eliminates T7’s susceptibility to PARIS, are positioned at the Ocr-AriA interface. Thus, these mutants’ effects likely arise not from an inability of the mutant Ocr proteins to dimerize, but rather from an inability to interact stably with AriA through the same interface. Ocr-AriA binding is further stabilized by several positively charged residues from both protomers of AriA (R212, R213, R244, R345, and R346), which line the interior of the CC interface and interact with the negatively charged surface of Ocr (Figure 4h). To test this idea, we designed two charge-reversal mutants in AriA – 2RE (R345E + R346E) and 4RE (R212E + R213E + R345E + R346E) – and tested these mutants’ ability to counter phage infection. We found that the R4E mutant completely lost the ability to defend against phage T7, suggesting that charge is a major determinant for the recognition of T7 Ocr (Figure 4j and Supplementary Figure 13c). We tested the same mutants against phage T4, which does not encode Ocr but does encode an unrelated DNA-mimicking anti-restriction protein, Arn^25^. While wild-type PARIS showed robust defense against phage T4 infection, the 4RE mutant showed no activity (Figure 4j and Supplementary Figure 13c). These data suggest that PARIS recognizes multiple phage-encoded factors through a common receptor pocket on its AriA subunit. Altogether, these data show that AriA is an infection sensor that binds and inhibits AriB in the absence of infection, then releases AriB in the presence of phage triggers.

### The PARIS effector AriB activates through homodimerization

While we were initially unable to express and purify soluble AriB, we found that incubating AriA-AriB with Ocr released soluble AriB for further analysis (Figure 4c). The elution position of AriB by size exclusion chromatography suggested that AriB might form an oligomeric complex. To measure the oligomeric state of AriB released from AriA, we performed mass photometry to examine the size exclusion chromatography fractions enriched for AriB in our release experiments. The mass photometry analysis revealed a dominant species with a calculated molecular weight of 68 kDa, closely matching the expected molecular weight of an AriB dimer (69.6 kDa; Figure 5a). As the amount of released AriB was too low for experimental structure determination, we predicted the structure of an AriB dimer using AlphaFold2^26^. The resulting models agreed closely with our cryoEM structure of the AriB monomer, and confidently predicted a dimer (ipTM score 0.92) (Figure 5b-c and Supplementary Figure 14a-c) assembled through a large interface spanning both domains. In the predicted structure, the two AriB active sites are positioned at opposite ends on one face of the dimer. The predicted dimer interface is highly conserved among AriB homologs (Figure 5d). To gain further support for this predicted structure, we re-examined 2D class averages from cryoEM analysis of AriA-AriB+Ocr. While most classes represented AriA-Ocr complexes (Figure 4), one 2D class closely resembled a 2D projection of the predicted AriB dimeric model (Figure 5e). While this dataset lacks sufficient particle numbers and distinct views of the putative AriB dimer to determine its structure, these data nonetheless lend support to the observation that AriB forms a homodimer once it is released from AriA.

**Figure 5.**
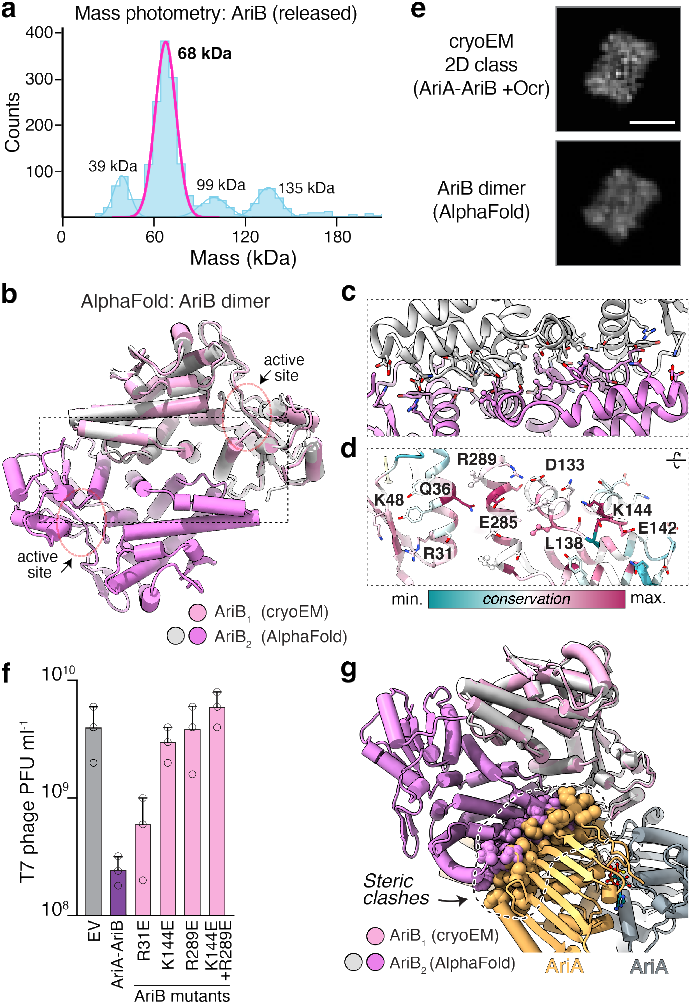
AriB forms a functional homodimer. **(a)** Mass photometry analysis of AriB released from AriA upon incubation with Ocr (see **Figure 4d**), revealing a predominant 68 kDa peak (1220 counts, 60% of total particles, σ 6.9 kDa) corresponding to the size of AriB dimer (theoretical molar mass 69.6 kDa). **(b)** AlphaFold 2 predicted structure of an AriB dimer (white/dark pink), superimposed on the monomeric AriB cryoEM structure (light pink). **(c)** Close-up view of the predicted AriB dimerization interface. Residues involved in dimerization are shown as sticks. **(d)** Conservation map (calculated by ConSurf^51^ from a sequence alignment of 30 AriB homologs) showing conservation of residues in the AriB dimerization interface. Highly conserved residues are colored magenta, and poorly conserved residues are colored cyan. **(e)** *Top:* A 2D class average from the cryoEM dataset collected after incubation of AriA-AriB^E90A^ with Ocr (see **Supplementary Figure 12**). *Bottom:* A simulated 2D projection of an AriB dimer, based on the predicted structure. Scale bar = 5 nm. **(f)** Analysis of EV (empty vector, from **Supplementary Figure 8e**), Wt (AriA-AriB system, cloned as an operon, from **Supplementary Figure 8e**), and AriB dimeric interface mutants for their ability to defend against T7 phage infection. R31E, K144E, R289E, K144E + R289E are the single and double charge-reversal mutants of AriB in AriA-AriB operon. Data represent the mean plaque-forming units (PFU) mL^−1^ of phage T7 from three independent replicates, with individual data points shown (n = 3; see representative plaque assay in **Supplementary Figure 14d**). **(g)** Modeling the predicted AriB dimer (white/dark pink) onto the AriB^E90A^ monomer in the AriA^EQ^-AriB^E90A^ complex (light pink) reveals significant steric clashes with AriA (gray/gold; clashing residues shown as spheres).

To better understand the functional relevance of AriB dimerization, we mutated conserved residues in the putative AriB dimerization interface. Charge-reversal mutations of interface residues K144 and R289 (K144E, R289E, and K144E + R289E) impaired the antiphage activity of PARIS (Figure 5f and Supplementary Figure 14d), suggesting that dimerization is required for the nuclease activity of AriB. To investigate how AriA restrains AriB dimerization and maintains it in an inactive state, we performed a structural superposition of the predicted AriB dimer model onto the AriB monomer within the AriA-AriB complex. This comparison revealed significant clashes between the dimerizing AriB protomer and the AriA protomer involved in the HH dimer interface (Figure 5b and 5g). Thus, AriA sequesters AriB in an inactive monomeric state, allowing AriB dimerization and activation only upon Ocr sensing.

### PARIS activation inhibits protein translation

Our findings above show that PARIS inhibits phage infection through cellular growth inhibition that relies on the expression of functional AriB. To gain insight into the underlying mechanism of AriB toxicity, we employed bacterial cytological profiling. This approach identifies the metabolic pathways targeted by drugs or other toxins through their effects on bacterial chromosomal condensation, cell shape, and overall cellular morphology^27^. Using *E. coli* MG1655 as an expression host, we overexpressed AriB, AriA-AriB, or AriA-AriB+Ocr, and performed fluorescence imaging of these cells. While the expression of AriA-AriB did not significantly alter the cytological profile of cells, cells overexpressing AriB or AriA-AriB+Ocr exhibited toroidal chromosomes and a widened cellular morphology (Figure 6a-c and Supplementary Figure 15a-c). These phenotypes are hallmarks of protein translation inhibitors including the ribosome-targeting drug chloramphenico^l27^ (Figure 6d and Supplementary Figure 15d). These results, in conjunction with AriB’s structural similarities to the M5 ribonuclease, suggest that PARIS activation inhibits cell growth by blocking protein translation, potentially through cleavage of as-yet unidentified essential cellular tRNAs or ribosomal RNAs.

**Figure 6.**
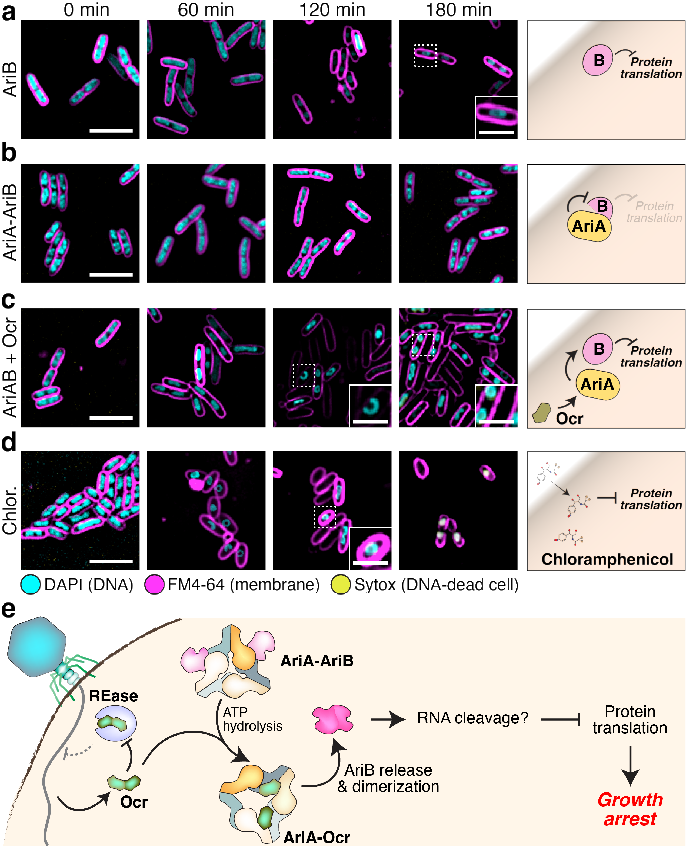
AriB inhibits protein synthesis. **(a-c)** *Left:* Fluorescence microscopy of *E. coli* MG1655 cells expressing AriB (panel **a**), AriA-AriB (panel **b**), or AriA-AriB + Ocr (panel **c**) at 0-, 60-, 120-, and 180-minutes post-induction of protein expression. Toroidal chromosomes indicate a global inhibition of protein translation^27^. Scale bars = 5 μm (2 μm for insets). In all images, cell membranes are stained with FM4-64 (magenta), DNA is stained with DAPI (cyan) for live cells, and SYTOX (yellow) for cells with permeabilized membranes. See **Supplementary Figure 15** for uncropped images. *Right:* Schematics illustrating findings from bacterial cytological profiling. **(d)** Fluorescence microscopy of *E. coli* MG1655 cells treated with chloramphenicol, a known inhibitor of protein translation. **(e)** Model for phage infection sensing and activation of the PARIS system.

## Discussion

Here we reveal the molecular mechanisms of the PARIS antiviral defense system, in which the AriA ATPase binds and inhibits the AriB nuclease in the absence of phage infection, then undergoes a significant conformational change coupled to ATP hydrolysis upon encountering the phage anti-restriction protein Ocr. This conformational change in AriA releases AriB and allows it to dimerize and become active. We hypothesize that, like other OLD-family nucleases that suppress global protein translation^28,29^, activated AriB cleaves one or more essential RNAs to block protein translation and suppress phage propagation. This mechanism is a striking example of how bacterial defense systems can synergize with one another for antiphage immunity: (1) first-responder defense systems target phage DNA; (2) the phage-encoded protein Ocr inhibits these systems; (3) the reserve defense system PARIS senses Ocr and halts cell growth (Figure 6e). Mutation of Ocr allows a phage to escape PARIS-mediated immunity, but these mutations could render the phage newly vulnerable to RM systems.

Our data show that selection pressure from both PARIS and the EcoKI-type RM system (encoded in *E. coli* MG1655) leads to the emergence of escaper phages capable of evading both defense systems. Prior work showed that a T7 phage carrying the Ocr F54V mutant escapes PARIS immunity while at the same time inhibiting EcoKI immunity^5^. PARIS escaper phages isolated in the absence of an active RM system showed a variety of mutations to Ocr including missense and nonsense mutations, which are expected to result in a complete loss of Ocr^5^. Thus, Ocr dimer interface mutants including F54V and L82R seem to represent a compromise: these Ocr mutants continue to inhibit RM-mediated immunity, but probably less effectively than wild-type Ocr. At the same time, both Ocr^F54V^ and Ocr^L82R^ are less prone to trigger PARIS immunity than wild-type Ocr (Supplementary Figure 10d)^5^. While our biochemical data shows that Ocr^L82R^ can cause limited AriB dissociation from AriA, Ocr^L82R^ fails to stably bind AriA and thereby likely allows AriA-AriB reassociation (Figure 4c). The low level of released AriB in the presence of Ocr^L82R^ may be inadequate to fully inhibit protein translation, giving the phage a window of opportunity to propagate. Overall, these data suggest that the malleability of Ocr enables phages to evade both bacterial defense and anti-anti-defense immune layers.

Rad50-like ABC ATPases and the related SMC ATPases play key roles in genome organization and maintenance in all cells^19^. These machines typically form homo- or heterodimers through two interfaces: one interface in the ATPase domains that is regulated by ATP binding and hydrolysis, and a second interface at the distal ends of their coiled-coil domains. Here we show that AriA, whose closest structural relatives are Rad50 and SMC ATPases, forms a distinctive heterohexamer by combining the canonical ATPase-domain dimer interface (here termed the HH interface) with a novel dimer interface involving its truncated coiled-coil domains (CC interface). The AriA HH interface serves as a docking site for AriB that is allosterically controlled both by binding of Ocr at the CC interface and by ATP hydrolysis. Thus, our data show how a Rad50-like ABC ATPase has evolved antiviral defense activity through a novel protein-protein binding surface and allosteric control of ATPase activity. Our data further suggest that the AriA CC interface recognizes multiple phage-encoded triggers, including T7 Ocr and potentially T4 Arn. We were unable to isolate T4 phage mutants that escape PARIS immunity in our *E. coli* strain encoding both PARIS and an EcoKI-type RM system (not shown), suggesting that mutations in Arn that could potentially escape PARIS immunity do not maintain the ability to inhibit RM immunity.

The broad family of toprim/OLD family nucleases was discovered over 50 years ago^30^, yet these enzymes’ regulatory mechanisms and biological roles remain mostly unknown. In particular, defense systems that include a toprim/OLD family nuclease and a Rad50/SMC-like ATPase are widespread^31^. These systems include the *Lactococcus* AbiL system^32^, an unnamed two-gene system identified in *Nostoc* sp.^8^, and the *Campylobacter* RloAB system^33^. Our sequence analysis and structure predictions (not shown) suggest that these systems are all structurally distinct from PARIS, hinting at surprising diversity within this family of defense systems. Among these ABC ATPase + toprim/OLD nuclease systems, the *Campylobacter* RloAB system is found within a type I RM system^33^. Given that PARIS is triggered by a phage protein that inhibits RM-mediated immunity, the identification of a PARIS-like defense system within an RM system supports the idea that these systems may synergize when fighting a phage infection. Further work will be needed to determine whether all ATPase + toprim/OLD systems function in antiphage defense, and to define these systems’ mechanisms of activation and phage protection.

Our data show that PARIS functions equivalently to a type II toxin-antitoxin (TA) system, with the antitoxin AriA binding and inhibiting the toxin AriB. While TA systems were first discovered and characterized as two-gene “addiction modules” that promote the maintenance of conjugative plasmids^34^, more recent work has revealed these systems’ profound importance for diverse physiological processes^35^, notably including antiviral defense^18^. These systems include retrons, in which a protein toxin is inhibited by a reverse transcriptase-ssDNA antitoxin complex^36^; the fused toxin-antitoxin protein CapRel^37^; the ToxIN protein-RNA TA system^38^; and others^18,39,40^. These systems sense the presence of diverse phage-encoded nucleic acids or proteins and respond by activating their respective toxin, which block key metabolic pathways and/or kill the host cell to inhibit phage replication. PARIS, and the larger class of ABC ATPase + toprim/OLD nuclease systems, reveals a new means for infection sensing and phage defense among TA systems.

## Acknowledgements

The authors thank Aaron Whiteley for sharing T4 and T7 phages, Marius Matyszewski and Yajie Gu for help with cryoEM data collection, Aaron Whiteley and Arshad Desai for critical reading of this manuscript, and members of the Corbett lab for helpful discussions. All electron microscopy data were collected at the UCSD CryoEM Facility, which was built and equipped with funds from UCSD and an initial gift from the Agouron Institute. The authors thank Farin Ahmed and Phillip Ordoukhanian at The Scripps Research Institute Biophysics and Biochemistry Core Facility for assistance with mass photometry. Q.L. is a recipient of a predoctoral fellowship from the American Heart Association. The authors acknowledge funding from the National Institutes of Health R35 GM144121 (to K.D.C.) and R01GM129245 (to J.P.) and the Howard Hughes Medical Institute Emerging Pathogens Initiative (to J.P. and K.D.C.). This paper was typeset with the bioRxiv word template by @Chrelli: www.github.com/chrelli/bioRxiv-word-template.

## Author contributions

Conceptualization: A.D. and K.D.C. Methodology: A.D., Q.L., J.P., and K.D.C. Validation: A.D., Q.L., E.E., and K.D.C. Formal analysis: A.D., Q.L., E.E., J.P., and K.D.C. Investigation: A.D., Q.L., and E.E. Data curation: A.D. and K.D.C. Writing, original draft: A.D. and K.D.C. Writing, review and editing: A.D., Q.L., E.E., J.P., and K.D.C. Visualization: A.D., Q.L., E.E., and K.D.C. Supervision: J.P. and K.D.C. Funding acquisition: J.P. and K.D.C.

## Data availability

The *E. coli* B185 AriA^EQ^ cryoEM reconstruction has been deposited at the Electron Microscopy Data Bank (EMDB; https://www.ebi.ac.uk/emdb/) under accession number EMD-42969. Corresponding coordinates have been deposited at the RCSB Protein Data Bank (RCSB PDB; http://www.rcsb.org) under accession number 8V49. The *E. coli* B185 AriA^EQ^-AriB^E90A^ complex (Form I) cryoEM reconstruction has been deposited at the EMDB under accession number EMD-42966. Corresponding coordinates have been deposited at the RCSB PDB under accession number 8V46. The *E. coli* B185 AriA^EQ^-AriB^E90A^ complex (Form II) cryoEM reconstruction has been deposited at the EMDB under accession number EMD-42967. Corresponding coordinates have been deposited at the RCSB PDB under accession number 8V47. The *E. coli* B185 AriA^EQ^-AriB^E90A^ complex (Form III) cryoEM reconstruction has been deposited at the EMDB under accession number EMD-42968. Corresponding coordinates have been deposited at the RCSB PDB under accession number 8V48. The AriA-Ocr complex cryoEM reconstruction has been deposited at the EMDB under accession number EMD-42965. Corresponding coordinates have been deposited at the RCSB PDB under accession number 8V45. All other data is available in supplementary information.

## Materials and Methods

### DNA cloning

For protein expression, *E. coli* B185 AriA (NCBI Accession # WP_001007866.1), AriB (WP_000093097.1), and T7 Ocr (NCBI Accession # NP_041954.1) genes were cloned into UC Berkeley Macrolab vectors 2-BT (Addgene #29666) and 13S-A (Addgene #48323) for N-terminal TEV protease-cleavable His_6_-tag and tagless constructs, respectively. For coexpression of AriA and AriB, the *ariAB* operon (reverse complement of bases 2,703,816-2,706,133 of the *E. coli* B185 genome, NCBI Accession # NZ_OU349838.1) was cloned into Macrolab vector 2-BT, resulting in the N-terminal TEV protease-cleavable His_6_-tagged AriA coexpressed with untagged AriB. PCR-based site-directed mutagenesis was performed to generate point mutants.

For toxicity assessment assays on solid media and bacterial growth curve experiments, AriB or AriA-AriB were cloned into an arabi-nose-inducible UC Macrolab vector 8-B (Addgene #37502; ampicillin-resistant) and Ocr was cloned into an IPTG-inducible pTrc99A-based vector (kanamycin-resistant). For plaque assays, Ari-AriB plus their native upstream and downstream flanking regions (reverse complement of bases 2,703,777-2,706,242 of NCBI Accession # NZ_OU349838.1) were cloned into vector pLOCO2 ^41^ in SbfI and NotI restriction sites.

### Protein expression and purification

Proteins were expressed in *E. coli* Rosetta2 pLysS (EMD Millipore) by growing cells at 37°C to an OD_600_ of 0.6-0.8, followed by induction with 0.33 mM IPTG. Cultures were incubated overnight at 20°C for the expression of protein/complexes. Following a 14-16 hour expression, cells were harvested by centrifugation, and the bacterial pellets were resuspended in ice-cold resuspension buffer containing 50 mM Tris pH 7.5, 300 mM NaCl, 10 mM imidazole, 10% glycerol, 2 mM β-mercaptoethanol. Resuspended cells were subjected to lysis using a sonicator, and the lysate was clarified using centrifugation. The proteins were purified using Ni^2+^ affinity chromatography (Ni-NTA Superflow, Qiagen). Eluted protein was concentrated and passed over a Superdex 200 Increase 10/300 GL size exclusion column (Cytiva) in a buffer containing 20 mM Tris pH 7.4, 250 mM NaCl, 2 mM β-mercaptoethanol.

His_6_-tagged T7 Ocr was expressed and purified as above. After Ni^2+^ affinity chromatography, the N-terminal His_6_-tag was cleaved by treating the purified protein with TEV protease, followed by tagless protein purification using anion-exchange chromatography (HiTrap Q HP, Cytiva) in a buffer with 20 mM Tris pH 7.5, 2 mM β-mercaptoethanol, and 50 mM to 1 M NaCl. Peak fractions were concentrated, and the sample was further purified to homogeneity using a Superdex 200 Increase 10/300 GL column (Cytiva) in a buffer containing 20 mM Tris pH 7.4, 250 mM NaCl, 2 mM β-mercaptoethanol. Purity was determined using SDS-PAGE analysis, and the samples were flash-frozen in liquid nitrogen and stored at -80°C until further use.

### Phage infection assay

Plaque-Forming Unit (PFU) estimations were used as a measure of phage infections. Briefly, LB agar plates were prepared with 100 μg/mL carbenicillin and left to dry for 2-3 days at room temperature. An overnight starter culture of *E. coli* MG1655 (harboring the pLOCO2 empty vector or pLOCO2 encoding PARIS (pLOCO2-*paris*)) was diluted to an OD_600_ of 0.05 in LB media and grown at 37°C until the OD_600_ reached 0.2. Next, 0.5 mL of this grown culture was mixed with 4.5 mL of preheated top agar (LB plus 0.35% agar), and the mixture was overlaid onto the plates. 10-fold dilutions of phage (T4 or T7) stocks were prepared in the phage buffer (40 mM Tris pH 7.5, 150 mM NaCl, 10 mM MgSO_4_, and 1 mM CaCl_2_) as the diluent. After 2 hours of air drying of plates, 5 μL volume of phage dilutions were spotted on these plates, followed by overnight incubation at room temperature (for phage T7) or at 37°C (for phage T4). The plaques were quantified the next day by counting individual plaques in the highest dilution that individual plaques are visible, and the reported values are the mean plus standard deviation of three biological replicates.

### Phage escaper generation and sequencing

To obtain T7 phage that escape PARIS immunity, we followed the approach described by Zhang et al^37^. In brief, 100 μL of 0.2 OD_600_ cultures harboring either an empty pLOCO2 or pLOCO2-*paris*, along with 10 mM MgSO_4_ and 1 mM CaCl_2_, were added to a sterile 96-well plate, as illustrated in Supplementary Figure 10a. Next, we performed 10-fold serial dilutions of T7 phage in the culture wells and incubated the plate at 30°C for 6-8 hours, and monitored the culture lysis. The lysed wells from the system carrying bacteria were pooled, and the next round of infection was performed until the cell death pattern between the control cells and cells carrying PARIS was similar. The resulting lysates were streaked on a lawn of bacteria to obtain single plaques. Purified phage from three individual plaques were obtained and subsequently sent for DNA sequencing to SeqCenter (Pittsburgh, Pennsylvania, USA).

### Toxicity assays

For toxicity assays on solid media, constructs cloned in arabi-nose-inducible vectors were transformed into *E. coli* MG1655 cells, and the transformants were selected on LB agar plates containing 100 μg/mL carbenicillin. For co-expression assays, an IPTG-inducible vector harboring the T7 *ocr* gene was co-transformed (alongside the arabinose-inducible constructs) into *E. coli* MG1655 cells, and the transformants were selected on LB agar plates containing both kanamycin (50 μg/mL) and carbenicillin (100 μg/mL). An individual colony was inoculated in LB broth containing the appropriate antibiotic(s), and cultures were allowed to grow overnight at 37°C in an incubator shaker. The next day, the cultures were back-diluted (OD_600_ ∼0.025) and further subjected to growth until the OD_600_ reached 0.25. 10-fold serial dilutions of these cultures were prepared in LB broth, and 7.5 μL of these dilutions were spotted onto LB agar plates containing the appropriate antibiotic and inducer (0.2% arabinose or 100 μM IPTG) or suppressor (0.4% glucose). The plates were incubated overnight at 37°C and were imaged the next day using a ChemiDoc Imaging System (Bio-Rad).

For bacterial growth curves, overnight cultures of *E. coli* MG1655 strains with plasmids of interest were grown at 37°C in LB plus appropriate antibiotics. Cultures were diluted OD_600_=0.1 in fresh LB with antibiotics and grown at 30°C until OD_600_=0.5. The cells were diluted back to OD_600_=0.1 with LB plus antibiotics and the inducers (0.2% or 0.5% arabinose plus 100 μM IPTG). 100 μL of diluted cultures were plated to standard clear 96 well plate with lid (Corning) and incubated in a plate reader (Tecan Sunrise) at 30°C. The OD_600_ of each well was measured every 5 minutes for 120 cycles (10 hours total) with medium intensity shaking. Each sample included three independent replicates on a single plate.

### Protein oligomeric state determination

For analysis of oligomeric states by size exclusion chromatography coupled to multi-angle light scattering (SEC-MALS), a 100 μL protein/complex samples (AriA, AriA-AriB, and Ocr) at 2 mg/mL was injected into a Superdex 200 Increase 10/300 GL column (Cytiva) in a buffer containing 20 mM Tris pH 7.4, 150 mM NaCl, 2 mM β-mercaptoethanol. Light scattering and refractive index profiles were collected by miniDAWN TREOS and Optilab T-rEX detectors (Wyatt Technology), respectively, and molecular weight was calculated using ASTRA v.8 software (Wyatt Technology).

For molecular mass determination by mass photometry, the released AriB sample (corresponding to the ∼14 mL peak on size exclusion chromatography performed using a Superdex 200 Increase 10/300 GL column) was diluted to 20 nM in a buffer containing 20 mM Tris, 150 mM KCl, and 1 mM DTT. Data acquisition and analysis were performed at the Biophysics and Biochemistry Core at Scripps Research in La Jolla, California, USA. The measurements were conducted using a Refeyn TwoMP mass photometer (Refeyn Ltd.) with a data acquisition time of 60 seconds. AcquireMP and DiscoverMP software were employed for data collection and analysis, respectively. Contrast-to-mass conversion was performed using a calibration with BSA (66 kDa) and thyroglobulin (330 kDa). The recorded events were fitted to Gaussian distributions to calculate the molecular mass and standard deviation of the Gaussian fit.

### ATPase activity measurement

ATP hydrolysis rates were measured using the ADP-Glo Kinase assay (Promega). Reactions were performed with 500 μM ultra-pure ATP in a reaction buffer containing 20 mM Tris pH 7.5, 150 mM NaCl, 1 mM DTT, and 10 mM MgCl_2_. 5 μL reactions containing 133 nM of protein/complexes were incubated at 37°C for 45 minutes. Reactions were quenched by adding ADP Glo reagent, followed by the kinase detection reagent. Luminescence was measured in a ProxiPlate-384 Plus (PerkinElmer) using a TECAN (Mannedorf, Switzerland) Infinite M1000 microplate reader. The quantification of hydrolyzed ATP was done by referencing a standard curve plotted with the known concentrations of ADP and ATP mixtures. The reported ATPase rate represents the mean values obtained from four technical replicates, with standard deviation error bars included.

### Interaction assay for AriA-AriB and Ocr

To investigate the interaction between AriA-AriB and Ocr, we employed size exclusion chromatography. Briefly, 6 μM (calculated as hetero-nonamer) AriA-AriB^E90A^ or AriA^EQ^-AriB^E90A^ were incubated 5 minutes at room temperature in a buffer containing 20 mM Tris, 150 mM KCl, 1 mM DTT, 10 μM ATP and 10 mM MgCl_2_. Next, 18 μM Ocr (calculated as a homodimer) was added, incubated 5-7 minutes at room temperature, then the mixture was cooled to 4°C. Samples were applied to a Superdex 200 Increase 10/300 GL column (Cytiva) in a buffer 20 mM Tris, 150 mM KCl, and 1 mM DTT. Fractions were separated on SDS-PAGE and visualized using Coomassie staining.

### CryoEM sample preparation and data acquisition

For the determination of AriA structure, we expressed His_6_-AriA^E393Q^ (hereafter termed AriA^EQ^), purified the protein through Ni^2+^ affinity chromatography, and performed, size exclusion chromatography in a buffer containing 20 mM Tris pH 7.4, 250 mM NaCl, 2 mM β-mercaptoethanol. The peak fraction was diluted to 7 μM AriA^EQ^ concentration (as a hexamer), and was mixed with 10 mM MgCl_2_ and 1 mM ATP. The sample mixture was then incubated on ice for 20 minutes. Prior to use, Quantifoil copper 1.2/1.3 300 mesh grids were plasma cleaned for 12 seconds using a preset program in the Solarus II plasma cleaner (Gatan). The AriA^EQ^ protein sample was applied to the grid in a 3.5 μL drop within the environmental chamber adjusted to 4°C temperature and approximately 95% humidity in a Vitrobot Mark IV (ThermoFisher Scientific). After a 5 seconds incubation, the grids were blotted with a blot force of 4 for 4 seconds, the sample was then plunged frozen into liquid nitrogen-cooled liquid ethane.

Grids were mounteds into standard AutoGrids (ThermoFisher Scientific) for imaging. Imaging was done using a Titan Krios G4 transmission electron microscope (ThermoFisher Scientific) operated at 300 kV configured for fringe-free illumination and equipped with a Falcon 4 direct electron detector with Selectris X energy filter. The microscope was operated in EFTEM mode with a slit-width of 20 eV. Automated data acquisition was performed using EPU (ThermoFisher Scientific). Movies were collected at a magnification of 130,000x and a pixel size of 0.935 Å, with a total dose of 50 e-/Å^2^ and a rate of 7 eps. Defocus range of -0.5 to -2.2 was used during the data collection. In total, 5,174 movies were used in the final data processing after applying the CTF fitting and excessive motion criteria.

For the determination of AriA^EQ^-AriB^E90A^ complex structure, we co-expressed His_6_-AriA^EQ^-AriB^E90A^ alongside tagless AriB^E90A^ on a separate vector to ensure fully-saturated AriA^EQ^, and purified the complex through Ni^2+^ affinity chromatography. Subsequently, size exclusion chromatography was performed in a buffer containing 20 mM Tris pH 7.4, 250 mM NaCl, 2 mM β-mercaptoethanol. The peak fraction containing the AriA-AriB complex was diluted to 6 μM concentration (as a hetero-nonamer, AriA_6_B_3_), mixed with 10 mM MgCl_2_ and 1 mM ATP, and incubated on ice for 20 minutes. Prior to use, UltrAuFoil 1.2/1.3 300 mesh grids were plasma cleaned for 12 seconds using a preset program in the Solarus II plasma cleaner (Gatan). The AriA^EQ^-AriB^E90A^ complex sample was applied to the grid in a 3 μL drop within the environmental chamber adjusted to 4°C temperature and approximately 95% humidity in a Vitrobot Mark IV (ThermoFisher Scientific). After a 4 second incubation, the grids were blotted with a blot force of 4 for 4 seconds, the sample was then plunged frozen into liquid nitrogen-cooled liquid ethane.

For imaging of the AriA^EQ^-AriB^E90A^ complex, data collection was performed using a Titan Krios G4 transmission electron microscope (ThermoFisher Scientific) as described above for AriA^E393Q^ imaging. In addition to the above specified settings, a 200 μm objective aperture was used. Defocus range of -0.8 to -2.2 was used during the data collection and 5,058 micrographs were included in the final data processing.

For AriA-Ocr structure determination, we separately expressed His_6_-AriA-AriB^E90A^ and Ocr and purified these proteins as described above. The N-terminal tag was removed using TEV protease and samples were further purified through size exclusion chromatography in a buffer containing 20 mM Tris pH 7.4, 150 mM KCl, 2 mM β-mercaptoethanol. Prior to use, UltrAuFoil 1.2/1.3 300 mesh grids were plasma cleaned for 12 seconds using a preset program in the Solarus II plasma cleaner (Gatan). Immediately prior to freezing samples, the AriA-AriB^E90A^-Ocr complex was prepared by diluting AriA-AriB^E90A^ to 6 μM (as a hetero-nonamer) in a buffer containing 20 mM Tris pH 7.4, 150 mM KCl, and 2 mM β-mercaptoethanol, 10 μM ATP, and 10 mM MgCl_2_. After incubation for 5 minutes at room temperature, Ocr was added to a final concentration of 18 μM (calculated as a homodimer). The mixture incubated at room temperature for 5-7 minutes, followed by cooling to 4°C. Immediately, the prepared complex was applied to the grid in a 3.5 μL drop within the environmental chamber adjusted to 4°C temperature and approximately 95% humidity in a Vitrobot Mark IV (ThermoFisher Scientific). After a 5 second incubation, the grids were blotted with a blot force of 4 for 4 seconds, and the sample was plunged frozen into liquid nitrogen-cooled liquid ethane.

Grids were mounteds into standard AutoGrids (ThermoFisher Scientific) for imaging. Imaging was done using a Titan Krios G4 transmission electron microscope (ThermoFisher Scientific) operated at 300 kV configured for fringe-free illumination and equipped with a Falcon 4 direct electron detector with Selectris X energy filter. The microscope was operated in EFTEM mode with a slit-width of 20 eV. Automated data acquisition was performed using EPU (ThermoFisher Scientific). Movies were collected at a magnification of 130,000x and a pixel size of 0.935 Å, with a total dose of 50 e-/Å^2^ and a rate of 7 eps. Defocus range of -0.8 to -2.2 was used during the data collection. In total, 6,821 movies were used in the final data processing after applying the CTF fitting, excessive motion criterias, and manual selection.

### CryoEM data processing and coordinate model building

All cryoEM datasets were processed using cryoSPARC version 4 ^42^. Movies were first motion-corrected using patch motion-correction (multi), and CTF estimations were performed using CTF estimation (multi). For the AriA^EQ^ sample, initial particle picking was executed using a blob-picking strategy with a diameter ranging from 120 to 300 Å. The selected particles were curated using 2D classification, and representative 2D classes were chosen for template-based particle picking from 5,174 micrographs. The resulting picks were manually inspected and subjected to multiple rounds of 2D classification. Approximately 301,000 curated particles were used for *ab-initio* 3D reconstruction in 4 different classes. Particle class re-assignment was performed using heterogeneous refinement. Ultimately, 106,506 particles were sorted and refined using the NU (non-uniform) refinement method in cryoSPARC with the “Fit Spherical Aberration,” “Fit Tetrafoil,” and “Fit Anisotropic Magnification” options enabled. This resulted in a map with a resolution of around 3.6 Å. For atomic model building, a local refinement was performed with a mask (encompassing two AriA chains interacting at their ATPase head interaction interface, and their coiled-coil interface regions with other interacting protomers) and a 3.45 Å resolution local refinement map was constructed. For the detailed data processing flowchart, please see Supplementary Figure 3.

For the AriA^EQ^-AriB^E90A^ complex, initial particle picking was executed using a blob-picking strategy with a diameter ranging from 120 to 300 Å. These particles were manually inspected and curated using the “Inspect Particle Picks” job in cryoSPARC. Multiple rounds of 2D classifications resulted in 574,500 particles which were used in template creation for template-based picking and Deep classification (see below). The template-based picking was followed by multiple rounds of 2D classifications, *ab-initio* reconstructions, heterogeneous refinement, and 3D classifications. 132,804 particles were used for the homogeneous reconstruction and NU refinement with “Fit Spherical Aberration”, “Fit Tetrafoil,” and “Fit Anisotropic Magnification” options enabled. This resulted in a map with a resolution of 3.09 Å. To enable model building, a local refinement was performed with a mask (encompassing AriA ATPase head and AriB EM-density region), and a 2.95 Å resolution local refinement map was constructed. For the detailed data processing flowchart, please see Supplementary Figure 6.

Based on the above processing and inspections of AriA^EQ^-AriB^E90A^ EM-maps, we observed that owing to its higher-order oligomerization, AriA subunits form three asymmetric interaction interfaces where AriB can bind. While this can result in nine polymorphic complex forms, we were unable to separate these forms using standard 3D classification jobs. Therefore, we used a knowledge-based modeling approach for the reconstructions of nine possible subunit configurations. These maps were included in the heterogeneous refinement jobs in cryoSPARC and particle subsets were separated in multiple classes. Finally, NU refinement was performed for all 9 particle classes. For the detailed data processing flowcharts and map class schematics, please see Supplementary Figure 9.

For the AriA-Ocr complex, initial particle picking was executed using a blob-picking strategy with a diameter ranging from 120 to 300 Å. We selected 2D classes that showed hexameric AriA-like features and generated an initial 3D map for template creation. Next, template-based picking was performed, particles were manually inspected, and the particle curation was performed through 2D classifications, multiple rounds of *ab-initio* 3D modeling, and heterogeneous refinement (see Supplementary Figure 12). Finally, we used 99,697 particles for map reconstruction, followed by NU refinement with the “Fit Spherical Aberration,” “Fit Tetrafoil,” and “Fit Anisotropic Magnification” options enabled. This workflow resulted in a 3.63 Å map, which was further subjected to a local refinement (3.46 Å) using a mask encompassing two AriA protomers involved in coiled-coil interactions, their extended ATPase head dimerizing subunit, and an Ocr monomer.

As with the AriA^EQ^-AriB^E90A^ sample, due to the non-correlated symmetry resulting from asymmetric Ocr binding to three AriA CC subunit structures built with the 3.46 Å map, we generated five structural templates representing these states and systematically sorted particles into the different classes using heterogeneous refinement. Independent processing of these datasets resulted in maps ranging from 3.68 Å to 9.17 Å in resolution (see Supplementary Figure 13a-b).

For atomic model building, initial monomer models were generated by AlphaFold2^26^. These models were manually docked into cryoEM maps in ChimeraX^43^ and rebuilt in COOT^44^, then subjected to real-space positional and B-factor refinement in phenix.refine^45^. For low resolution regions, subunit structures built into high-resolution local refinement maps were used as reference models for reference-based refinement in phenix.refine^45^.

### Bacterial cytological profiling

Bacterial cytological profiling was performed as described by Nonejuie et al^27^. In brief, arabinose- or IPTG-inducible constructs expressing proteins of interest were transformed into *E. coli* MG1655 cells. The resulting transformants were grown at 37°C in LB broth plus appropriate antibiotics overnight. The next day, cultures were diluted to OD_600_ ∼0.1 in fresh LB media with antibiotics and grown at 30°C until the OD_600_ reached 0.2. Cultures were then supplemented with inducers (0.2% arabinose and 100 μM IPTG) and grown at 37°C. Samples were collected at 0 MPI (minutes post induction), 60 MPI, 120 MPI, and 180 MPI. For each sample, 20 μL of bacterial cell suspension was mixed with 1 μL dye mix (1 μL of 1 mg/mL FM4-64, 2 μL of 0.25 mM SYTOX Green, 1 μL of 2 mg/mL DAPI) in a 1.7 mL Eppendorf tube. 10 μL of the suspension was spotted on an agarose-pad slide (1% agarose with 25% LB). After 5 minutes of incubation at room temperature, a glass coverslip was placed on each pad slide and samples were imaged with fluorescence microscopy (Deltavision Elite System, Cytiva).

### Protein structure prediction using AlphaFold2

For AriB dimer model prediction, we used AlphaFold2^26^ as implemented by the ColabFold project^46^. MMseqs2 and AMBER force field options were used for multiple sequence alignment and model relaxation, respectively^47,48^.

### Computational analysis and figure generation

Structural homologs were identified using the DALI server^49^. Figures were prepared with ChimeraX^43^. Fluorescence microscopy images were processed and visualized using Fiji^50^. Graphpad Prism version 10 was used for graph generation and statistical analysis.

## Supplementary Material

Supplementary material is available online at bioRxiv.

