## Supplementary Data for "Architecture and infection-sensing mechanism of the bacterial PARIS defense system"

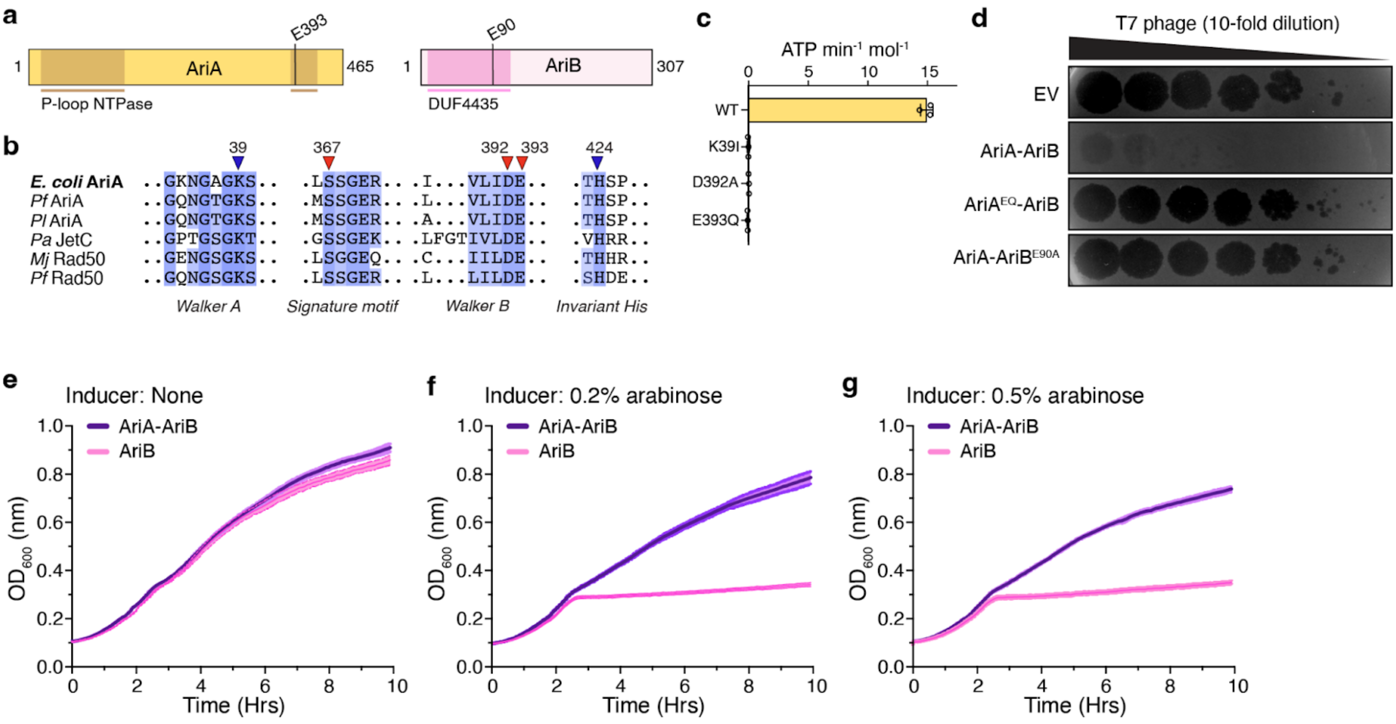


**Supplementary Figure 1 | PARIS is an antiphage toxin-antitoxin system. (a)** Schematic representation of the domains in the *E. coli* B185 AriA and AriB subunits, with predicted domain annotations. **(b)** Multiple sequence alignment showing conservation of amino acid residues involved in ATP binding and catalysis in AriA and Rad50 homologs (*E. coli* AriA: NCBI accession # WP_001007866.1; *P. sp* FW300-N1A1 AriA: NCBI accession # WP_103399253.1; *P. luteoviolacea* AriA: NCBI accession # WP_063361701.1; *P. aeruginosa* PA14 JetC: NCBI accession # WP_016254338.1; *M. jannaschii* Rad50: NCBI accession # Q58718.1; *P. furiosus* Rad50: NCBI accession # WP_011012307.1). The conserved residues involved in catalysis are indicated. **(c)** ATPase activity of AriA and its conserved residue mutants. K39I: a Walker-A motif mutant designed to disrupt ATP binding. D392A: a Walker-B motif mutant designed to disrupt ATP binding. E393Q: a Walker-B motif mutant designed to disrupt ATPase activity. ATP hydrolysis is expressed as moles of ATP hydrolyzed per minute per mole of AriA hexamer. Error bars represent the average and standard deviation of three measurements (n = 3; open circles). **(d)** A representative plaque-forming unit assay, demonstrating T7 phage plaques on the *E. coli* MG1655 bacterial lawn carrying either an empty vector (EV), the PARIS system (AriA-AriB, as an operon), the ATPase mutant AriA$^{E393Q}$ (AriA$^{EQ}$-AriB), or a putative nuclease-dead mutant AriB$^{E90A}$ (AriA-AriB$^{E90A}$). T7 phage 10-fold dilutions are spotted as indicated with a gradient. **(e, f, g)** Growth curve assays depicting the effects of AriB or AriA-AriB overexpression in *E. coli* MG1655 bacterial cells. The presence and absence of inducers, as well as their relative concentrations, are indicated above. Curves and error bars represent average +/- standard deviation from three independent replicates (n = 3), and are representative of two independent trials.

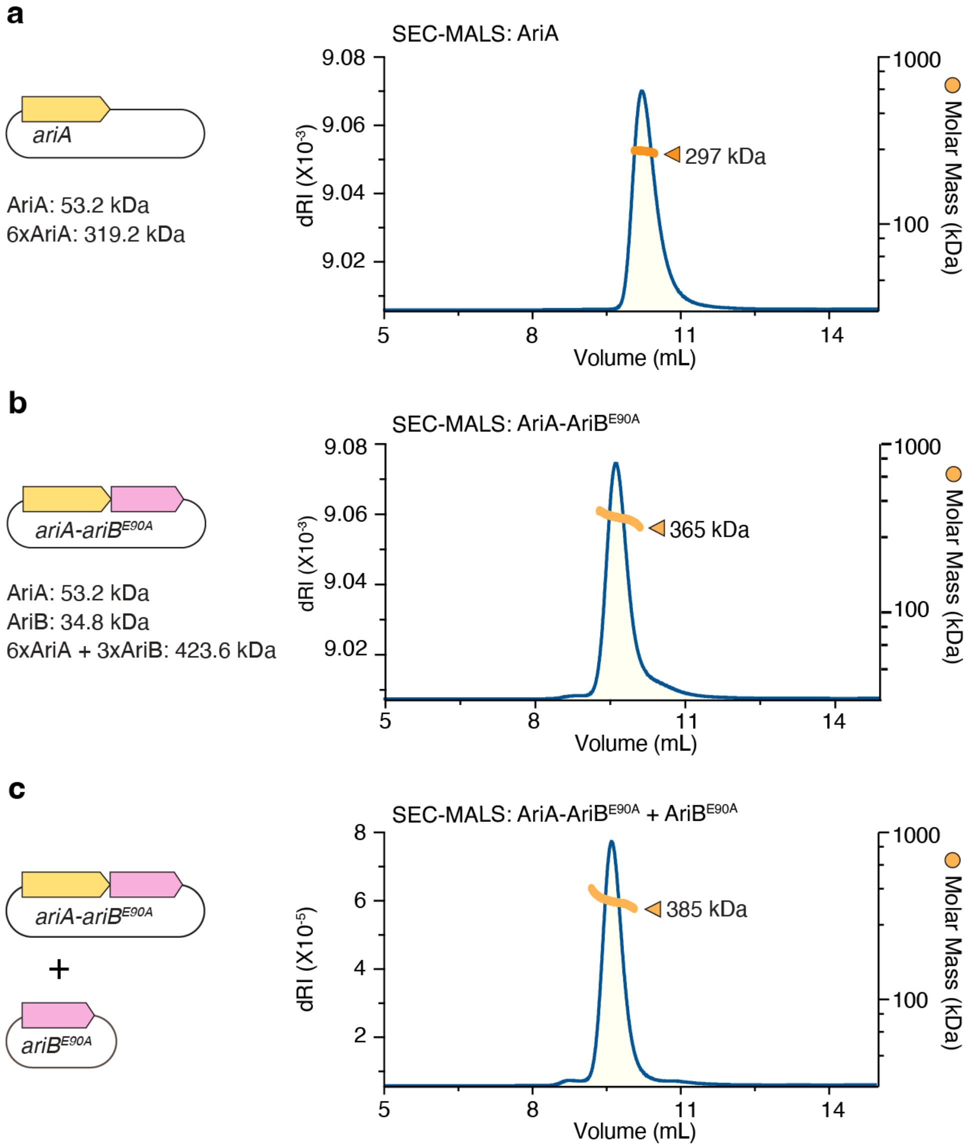


**Supplementary Figure 2 | Oligomeric states of AriA and AriA-AriB complex. (a, b, c)** Schematic representations illustrating expression strategies and size exclusion chromatography coupled to multi-angle light scattering (SEC-MALS) analysis of AriA, AriA-AriB$^{E90A}$, and AriA-AriB$^{E90A}$ + AriB$^{E90A}$ complexes. Blue lines depict protein concentration (dRI; change in refractive index from baseline), while orange lines indicate measured molecular mass in kDa. The theoretical molecular weight for each subunit and the inferred complexes are indicated beneath the expression strategy scheme.

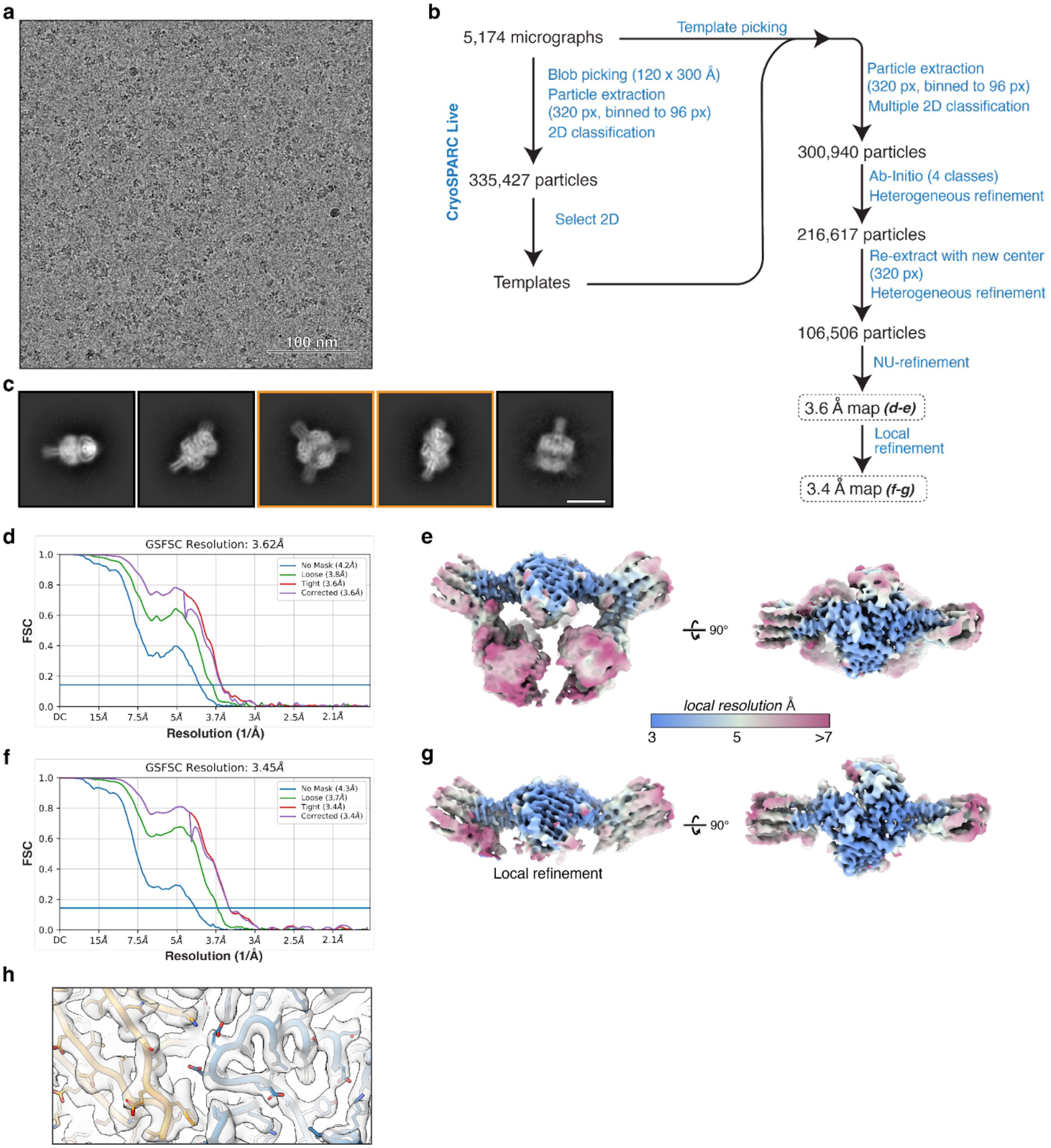


**Supplementary Figure 3 | CryoEM workflow for AriA$^{EQ}$.** **(a)** Representative micrograph displaying the AriA$^{EQ}$ sample. Scale bar = 100 nm. **(b)** Workflow outlining the cryoEM reconstruction process for AriA$^{EQ}$ using cryoSPARC. **(c)** Representative 2D classes displaying AriA samples from the final particle stack, with the highlighted yellow classes also depicted in **Figure 2b**. Scale bar = 10 nm. **(d)** Gold-standard FSC curve for the final global refinement of AriA$^{EQ}$. **(e)** Two views of the globally refined cryoEM map for the AriA$^{EQ}$ hexamer, color-coded by local resolution from <3 Å (blue) to >7 Å (red). **(f)** Gold-standard FSC curve for the masked local refinement of AriA$^{EQ}$. **(g)** Two views of the locally refined cryoEM map for the AriA$^{EQ}$ hexamer, color-coded by local resolution from <3 Å (blue) to >7 Å (red). **(h)** Example cryoEM density with a built atomic model demonstrating the model fit. Yellow and blue are two AriA$^{EQ}$ protomers.

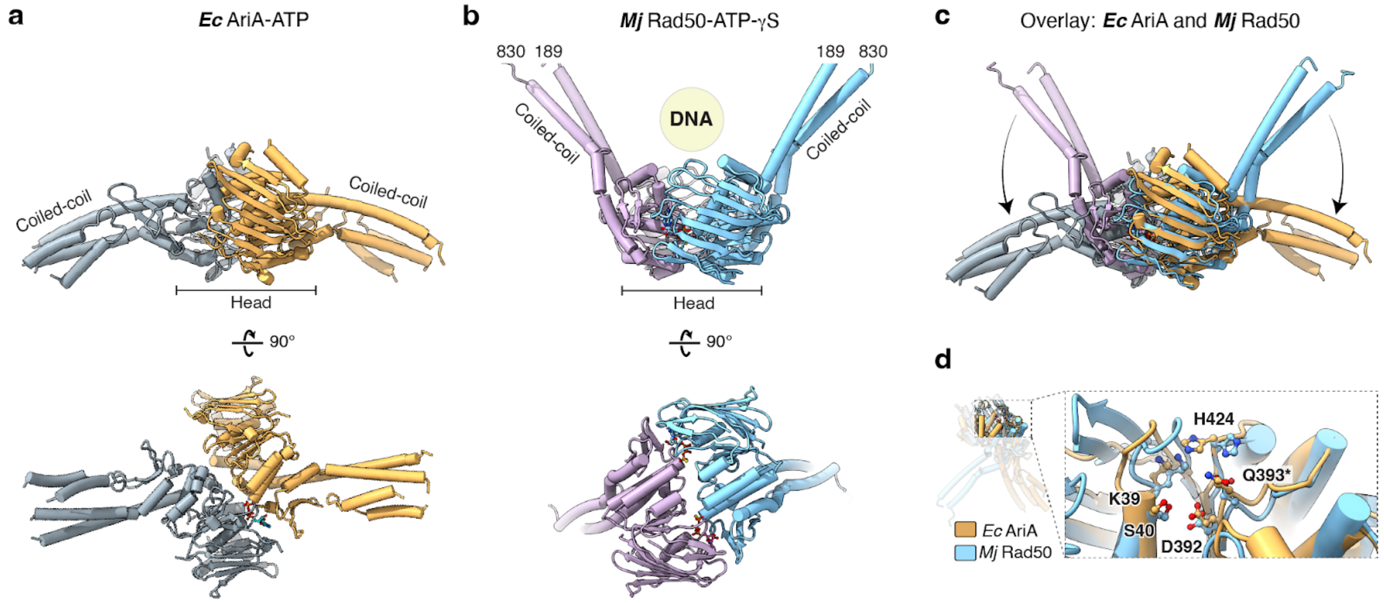


**Supplementary Figure 4 | Comparative analysis of *E. coli* B185 AriA sensory subunit and its structural homolog. (a)** Two views of the AriA HH dimer, with ATPase heads and coiled coils labeled. **(b)** Two views of the *M. jannaschii* Rad50 dimer (PDB ID 3AV0; no associated publication) equivalent to the AriA views in panel a. **(c)** Overlay of AriA and Rad50 homodimer depicting the distinct coiled-coil orientation. **(d)** Closeup view of the AriA and *Mj* Rad50 ATPase active site, with conserved AriA active site residues shown as sticks and labeled (see multiple sequence alignment in **Supplementary Figure 1b**). Q393* indicates the E393Q mutation.

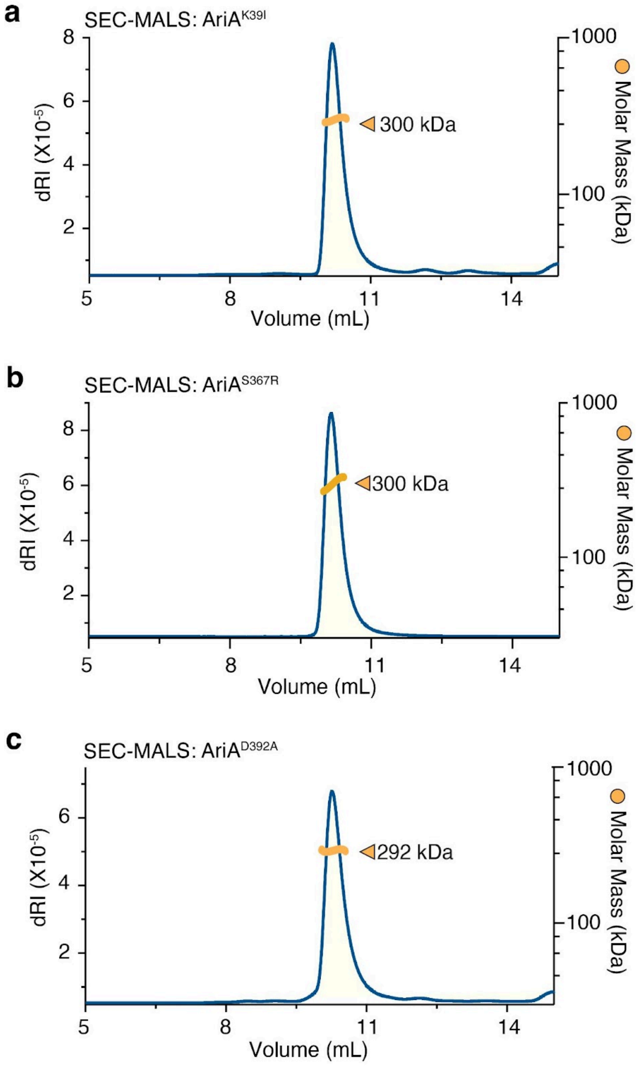


**Supplementary Figure 5 | Characterization of AriA ATPase domain mutants. (a, b, c)** Size-exclusion chromatography coupled to multi-angle light scattering (SEC-MALS) analysis of AriA$^{K39I}$, AriA$^{D392A}$, and AriA$^{S367R}$ (see **Supplementary Figure 2a** for SEC-MALS analysis of wild-type AriA). K39I: a Walker A motif mutant designed to disrupt ATP binding. D392A: a Walker B motif mutant designed to disrupt ATP binding. S367R: a signature motif mutant designed to disrupt ATPase head-head interaction interface. Blue lines depict protein concentration (measured as a change in refractive index, dRI), while orange lines indicate measured molecular weight (molar mass in kDa). The theoretical molecular weight for an AriA monomer is 53.2 kDa, and for a hexamer is 319.2 kDa.

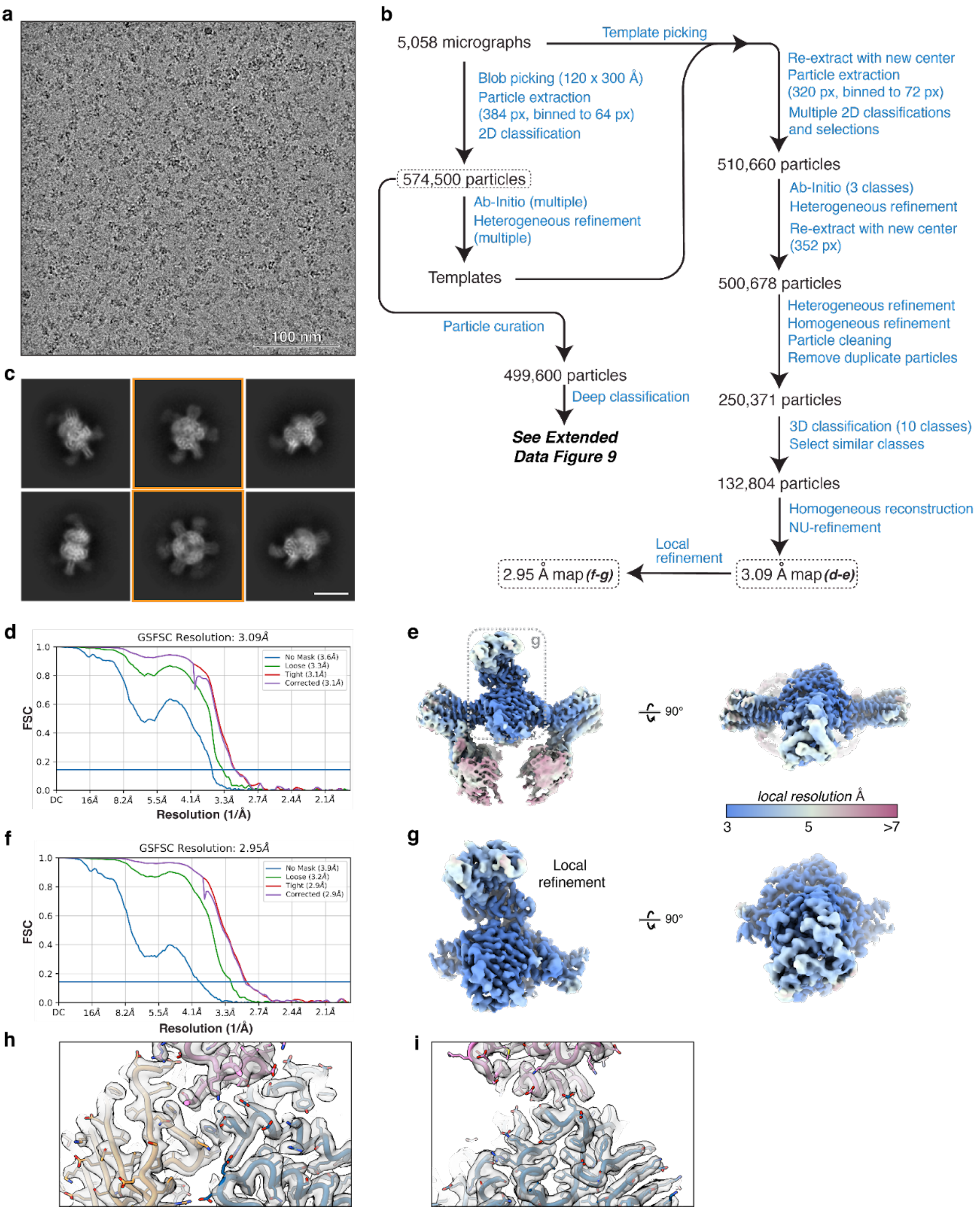


**Supplementary Figure 6 | CryoEM Workflow for AriA$^{EQ}$-AriB$^{E90A}$. (a)** Representative micrograph for the AriA$^{EQ}$-AriB$^{E90A}$ complex. Scale bar = 100 nm. **(b)** Workflow outlining the cryoEM reconstruction process for *E. coli* AriA$^{EQ}$-AriB$^{E90A}$ complex using cryoSPARC. **(c)** Representative 2D classes displaying AriA$^{EQ}$-AriB$^{E90A}$ samples from the final particle stack, with the highlighted yellow classes also depicted in **Figure 3b**. Scale bar = 10 nm. **(d)** Gold-standard FSC curve illustrating the final global refinement of AriA$^{EQ}$-AriB$^{E90A}$. **(e)** Two views presenting the globally refined cryoEM map for AriA$^{EQ}$-AriB$^{E90A}$, color-coded by local resolution from <3 Å (blue) to >7 Å (red). **(f)** Gold-standard FSC curve for the masked local refinement of AriA$^{EQ}$-AriB$^{E90A}$. **(g)** Two views showcasing the locally refined cryoEM map for AriA$^{EQ}$-AriB$^{E90A}$, color-coded by local resolution from <3 Å (blue) to >7 Å (red). **(h, i)** Example cryoEM density with a built atomic model demonstrating the model fit. Yellow and blue are two AriA protomer chains, and pink corresponds to AriB.

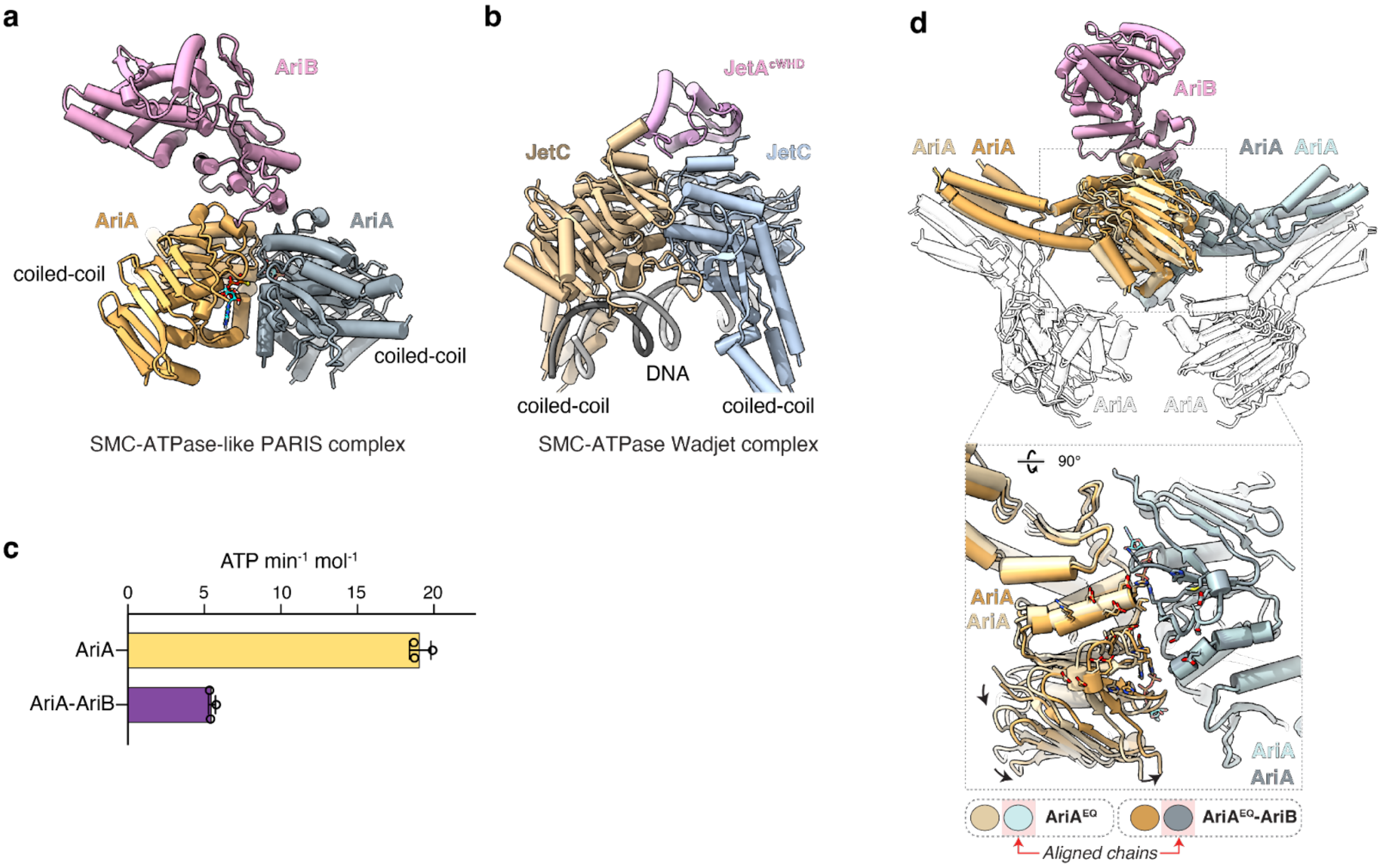


**Supplementary Figure 7 | Structural and functional changes in AriA upon interaction with AriB. (a, b)** Two similarly oriented views of AriA$^{EQ}$-AriB$^{E90A}$ and *Pseudomonas aeruginosa* PA14 Wadjet complex (PDB ID 8DK3)[20], illustrating that the AriB binding interface is canonically conserved among other SMC ATPases. **(c)** ATPase activity of AriA and AriA-AriB complex. ATP hydrolysis is expressed as moles of ATP hydrolyzed per minute per mole of AriA (as $AriA_6$) or AriA-AriB (as $AriA_6B_2$). Error bars represent the average and standard deviation of three independent measurements (n = 3; open circles). **(d)** Structural overlay presenting AriA$^{EQ}$ and AriA$^{EQ}$-AriB$^{E90A}$ complexes. The two molecules are overlaid by aligning two AriA protomer chains shown in light-green/gray colors. At the bottom, a close-up view depicts changes in the AriA ATPase head region upon binding with AriB. The residues involved in AriB binding are shown in stick representation for side chains. AriB is omitted for clarity.

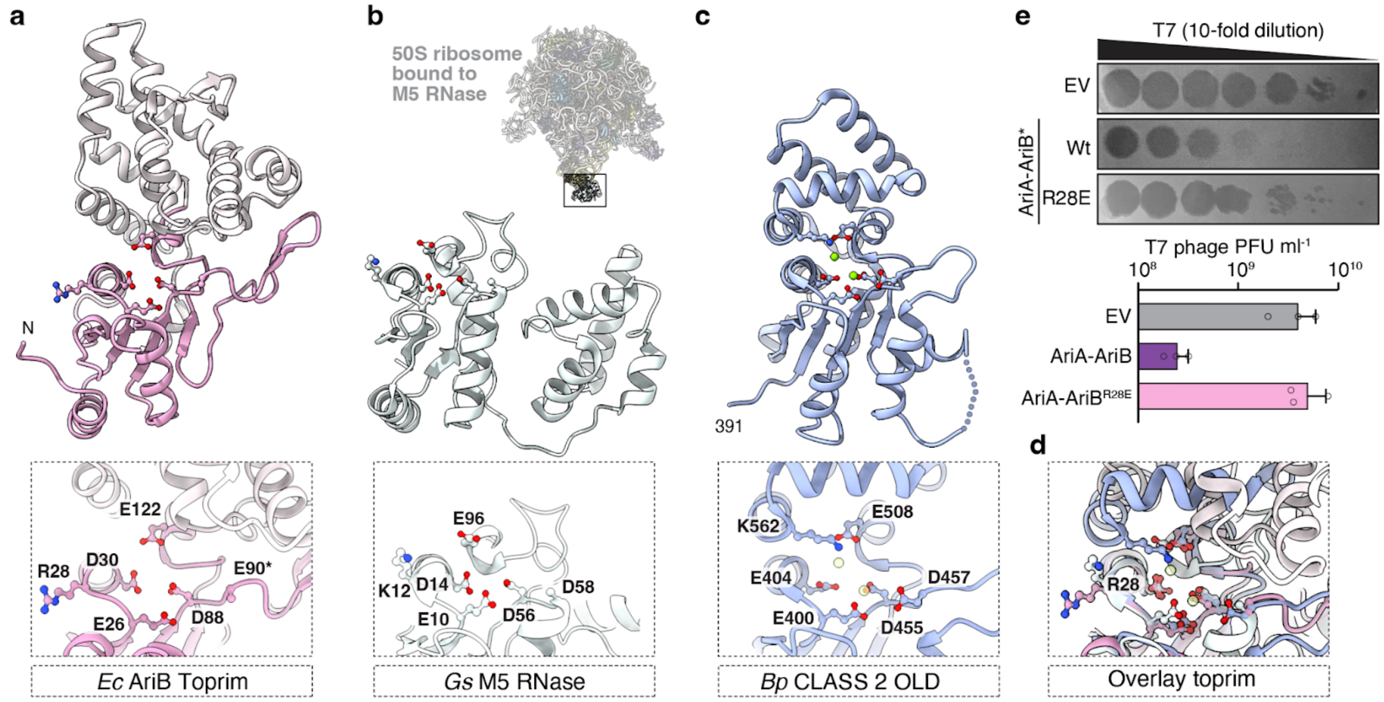


**Supplementary Figure 8 | Structural comparisons of AriB and its nuclease homologs. (a)** Cartoon representation of AriB monomer, with the toprim domain in pink and helical bundle domain in white. At the *Bottom;* Close-up view of the AriB active site. **(b)** Overall structure of *Geobacillus stearothermophilus* (*Gs*) M5 RNase bound to the 50S ribosome. M5 RNase is shown in a contrast clear window, zoomed in cartoon view below (PDB ID 6TPQ)[24]. *Bottom:* Close-up view of the active site of M5 RNase. **(c)** Cartoon structure of OLD DNase from *Burkholderia pseudomallei* (PDB ID 6NK8)[52]. Two bound metal ions are shown as green spheres. *Bottom:* Close-up view of the active site. **(d)** Overlay of active site residues from AriB, M5 RNase, and OLD DNase. The active site residues are shown in ball-and-stick representation. **(e)** *Top:* A representative plaque-forming unit assay, demonstrating T7 phage plaques on the *E. coli* MG1655 bacterial lawn carrying either an empty vector (EV), the PARIS system (Wt; AriA-AriB as an operon), or the RNA binding mutant AriB$^{R28E}$ (AriA-AriB$^{R28E}$). T7 phage 10-fold dilutions are spotted as indicated with a gradient. *Bottom:* Data represent the mean plaque-forming units (PFU) mL$^{-1}$ of phage T7 from three independent replicates, with individual data points shown (n = 3).

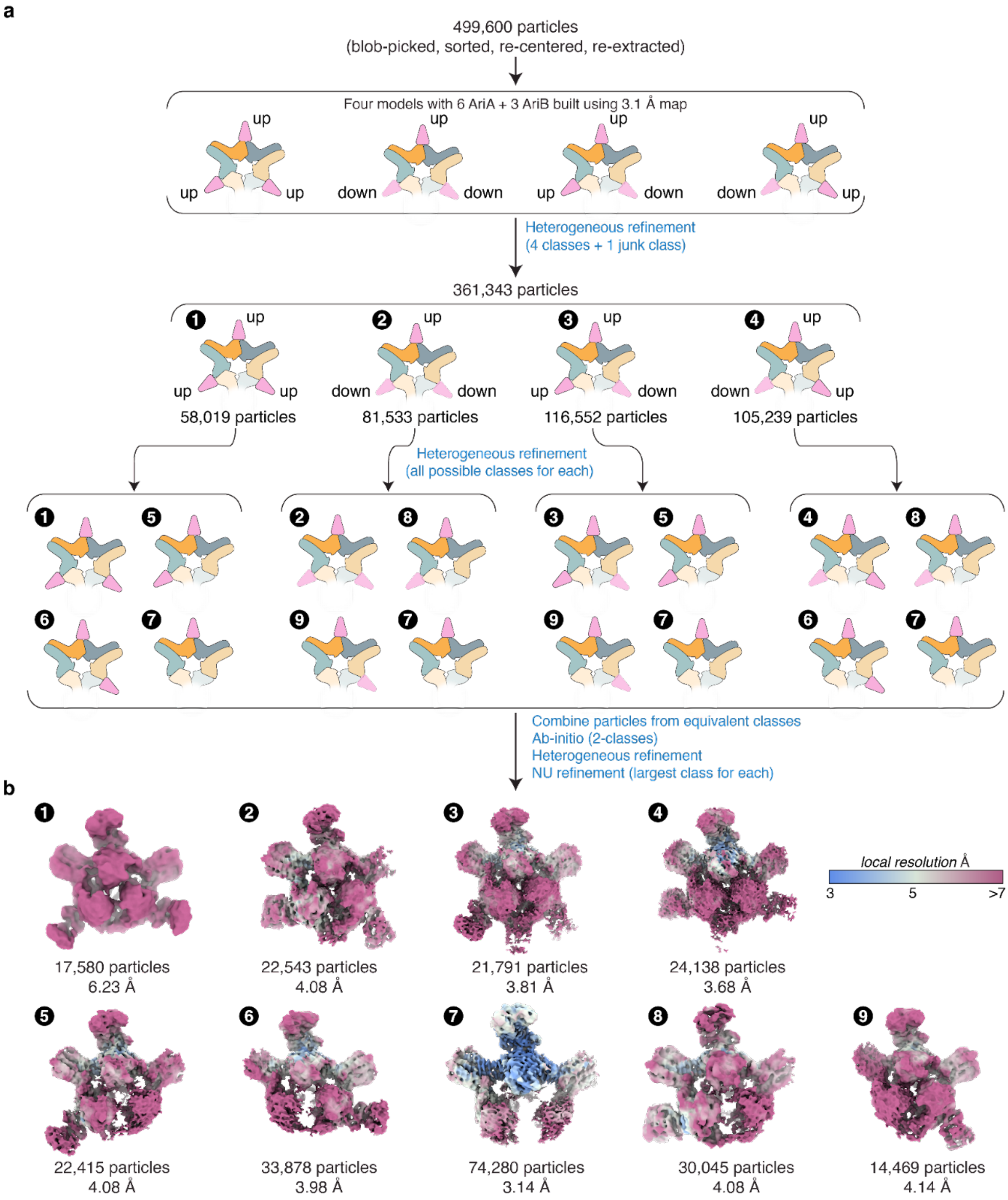


**Supplementary Figure 9 | CryoEM data processing workflow used in sorting polymorphic AriA$^{EQ}$-AriB$^{E90A}$ complexes. (a)** Systematic sorting of compositionally and conformationally heterogeneous AriA$^{EQ}$-AriB$^{E90A}$ complexes. The cartoon schematics show the possible arrangements of AriA$^{EQ}$ and AriB$^{E90A}$ subunits. Volume templates were generated using subunit structures built from the maps shown in **Supplementary Figure 6**. These volumes were imported into cryoSPARC and included in the "Heterogeneous refinement" jobs for particle class reassignment. **(b)** The final nine cryoEM reconstructions for the AriA$^{EQ}$-AriB$^{E90A}$ complex, color-coded by local resolution from <3 Å (blue) to >7 Å (red). The final resolution and the number of particles corresponding to each complex form are noted at the bottom of each map.

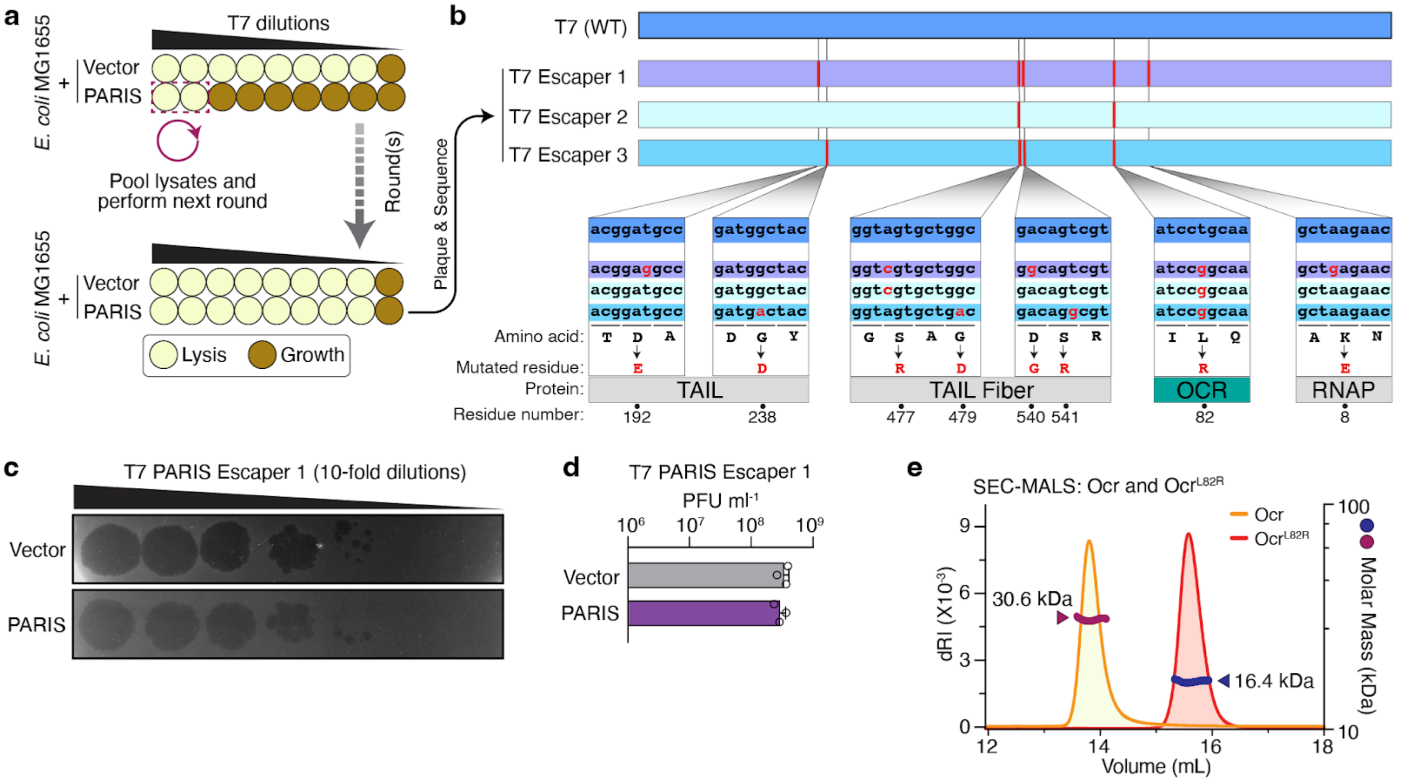


**Supplementary Figure 10 | Identification of T7 phage molecular pattern recognized by *E. coli* B185 PARIS system. (a)** Overview of the experimental evolution strategy used to obtain T7 phage escaper mutants against PARIS[37]. **(b)** Summary of the mutations identified in three T7 phage escaper mutants. Close-up views below show the DNA sequence alignment and the impact of mutations in the noted phage genes. **(c)** Representative plaque-forming unit assay demonstrating T7 escaper phage #1 plaques on an *E. coli* MG1655 bacterial lawn carrying either an empty vector (Vector) or the PARIS system. **(d)** Analysis of the empty vector (Vector) and PARIS system for their ability to defend against T7 PARIS escaper phage #1. The data represent mean and standard deviation of plaque-forming units (PFU) mL$^{-1}$ from three independent replicates, with individual data points shown as open circles (n = 3). **(e)** SEC-MALS analysis of T7 Ocr (orange curve with red molar mass measurement) and Ocr$^{L82R}$ (red curve with blue molar mass measurement). Theoretical molar mass of His$_6$-tagged Ocr = 16.1 kDa (monomer)/32.2 kDa (dimer).

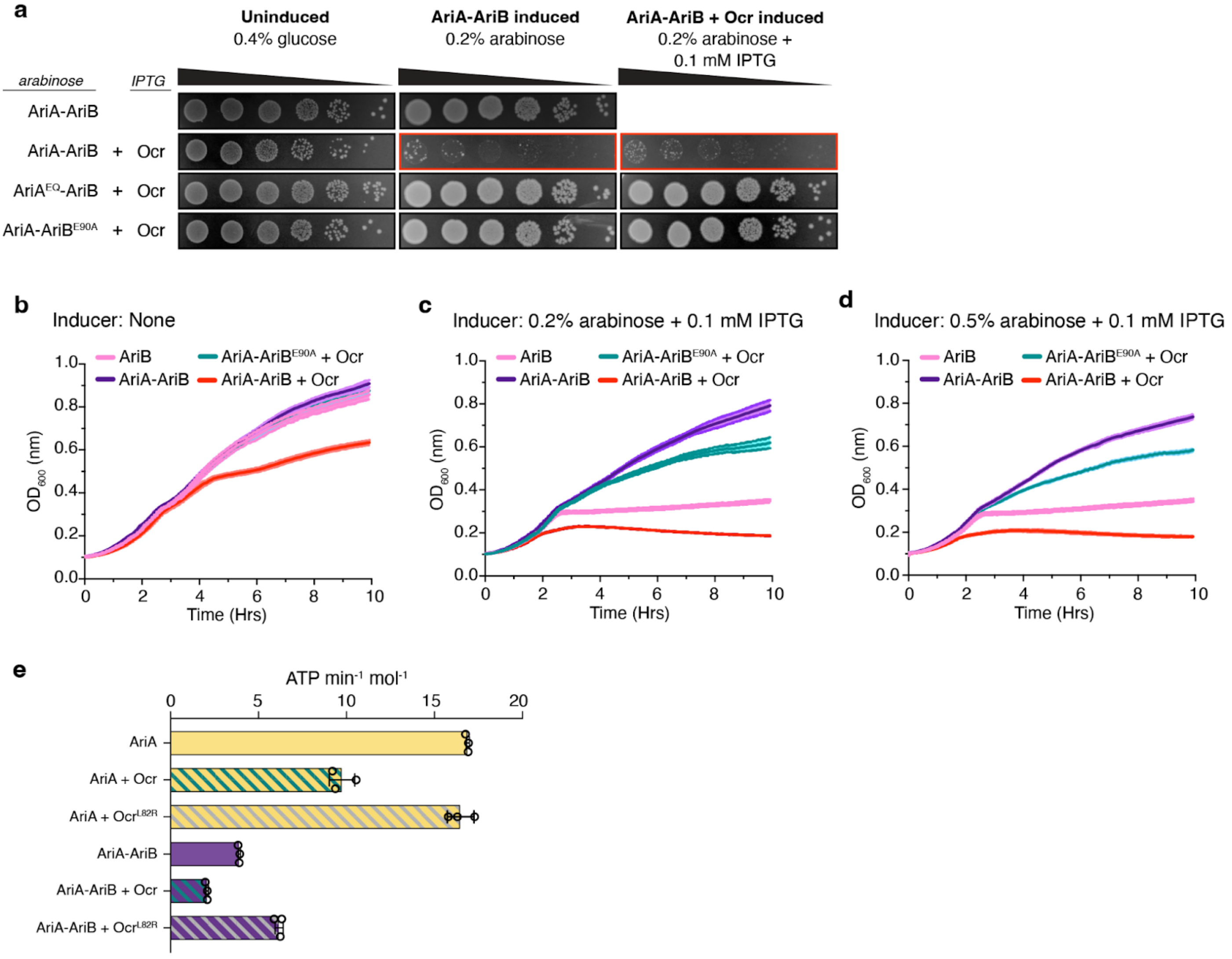


**Supplementary Figure 11 | Ocr triggers PARIS and inhibits ATP hydrolysis by AriA. (a)** Bacterial dilution spotting (10-fold) assay performed to measure the toxicity of AriA-AriB (from **Figure 1c**), AriA$^{EQ}$-AriB, and AriA-AriB$^{E90A}$ upon co-expression with Ocr. *Uninduced:* LB media + 0.4% glucose (suppressor) + antibiotics; *AriA-AriB Induced:* LB media + antibiotics + 0.2% arabinose. *AriA-AriB + Ocr Induced:* LB media + antibiotics + 0.2% arabinose + 0.1 mM IPTG. Cell growth inhibition observed when coexpressing AriA-AriB plus Ocr without Ocr induction (red outline) is likely due to low-level leaky expression of Ocr. **(b, c, d)** Growth curves of *E. coli* MG1655 cells transformed with plasmids carrying AriB (pink; from **Supplementary Figure 1d-f**), AriA-AriB (purple, from **Supplementary Figure 1d-f**), AriA-AriB + T7 Ocr (red), or AriA-AriB$^{E90A}$ (green), in the absence (panel b) or the presence (panels c-d) of inducers. Curves represent the average and standard deviation from three independent replicates (n = 3). **(e)** ATPase activity assay showing the effect of Ocr and Ocr$^{L82R}$ on AriA or AriA-AriB complex ATPase activity. ATP hydrolysis is expressed as moles of ATP hydrolyzed per minute per mole of AriA hexamer. Error bars represent the average and standard deviation of three independent measurements (n = 3; open circles).

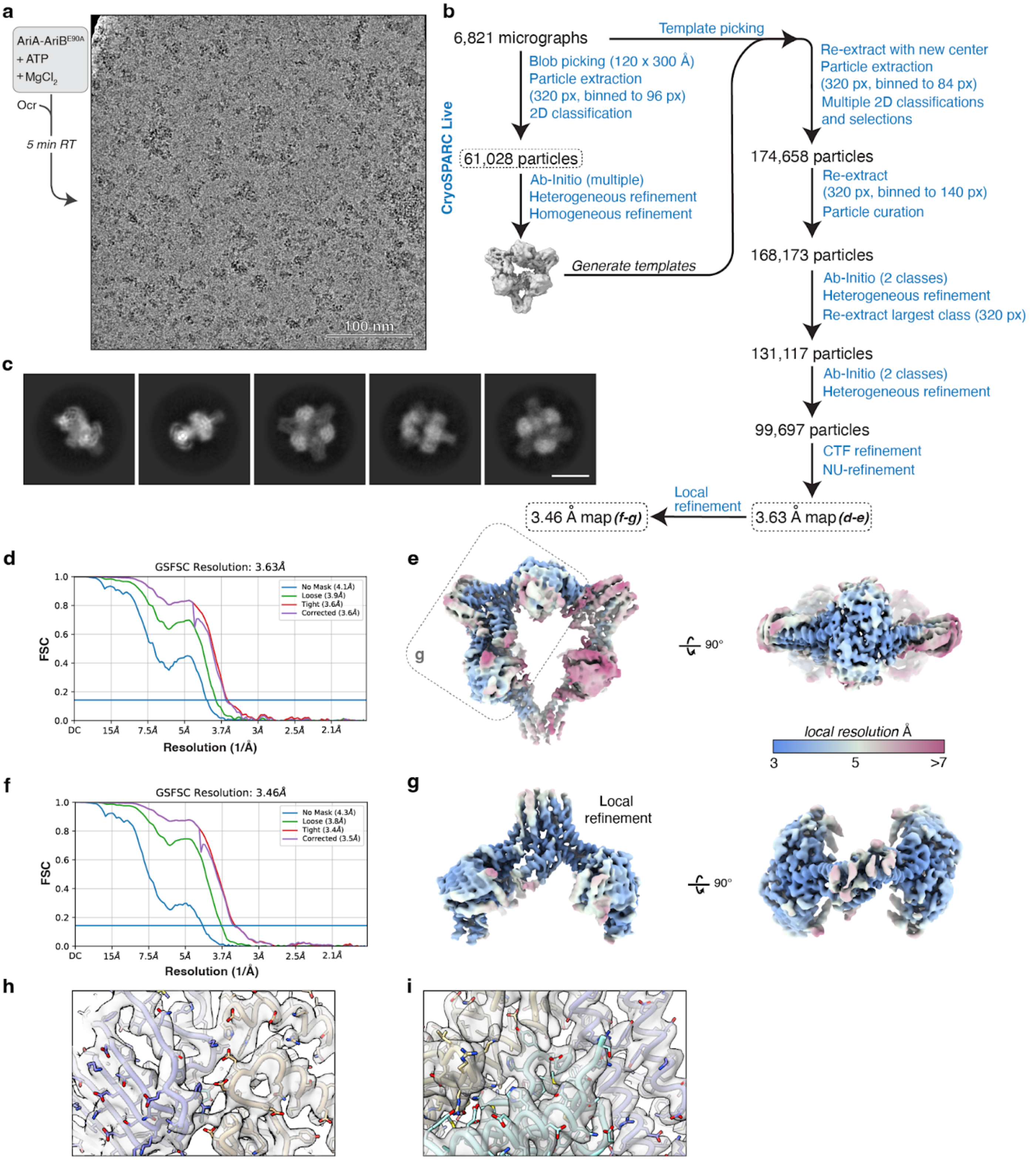


**Supplementary Figure 12 | CryoEM workflow for the AriA-Ocr complex. (a)** Sample preparation strategy and a representative micrograph displaying the PARIS + Ocr sample. Scale bar = 100 nm. **(b)** Workflow outlining the cryoEM data processing for the AriA-Ocr complex using cryoSPARC software. **(c)** Representative 2D classes showing AriA-Ocr samples from the final particle stack. Scale bar = 10 nm. **(d)** Gold-standard FSC curve illustrating the final global refinement of the AriA-Ocr complex reconstruction. **(e)** Two views of the globally-refined cryoEM map for the AriA-Ocr complex, color-coded by local resolution from <3 Å (blue) to >7 Å (red). **(f)** Gold-standard FSC curve for the masked (noted in panel e) local refinement of the AriA-Ocr interaction interface region. **(g)** Two views of the locally refined cryoEM map for the AriA-Ocr interaction interface region, color-coded by local resolution from <3 Å (blue) to >7 Å (red). **(h, i)** Example cryoEM density with a docked atomic model demonstrating the model fit. Yellow and blue represent two AriA protomers, and green represents Ocr.

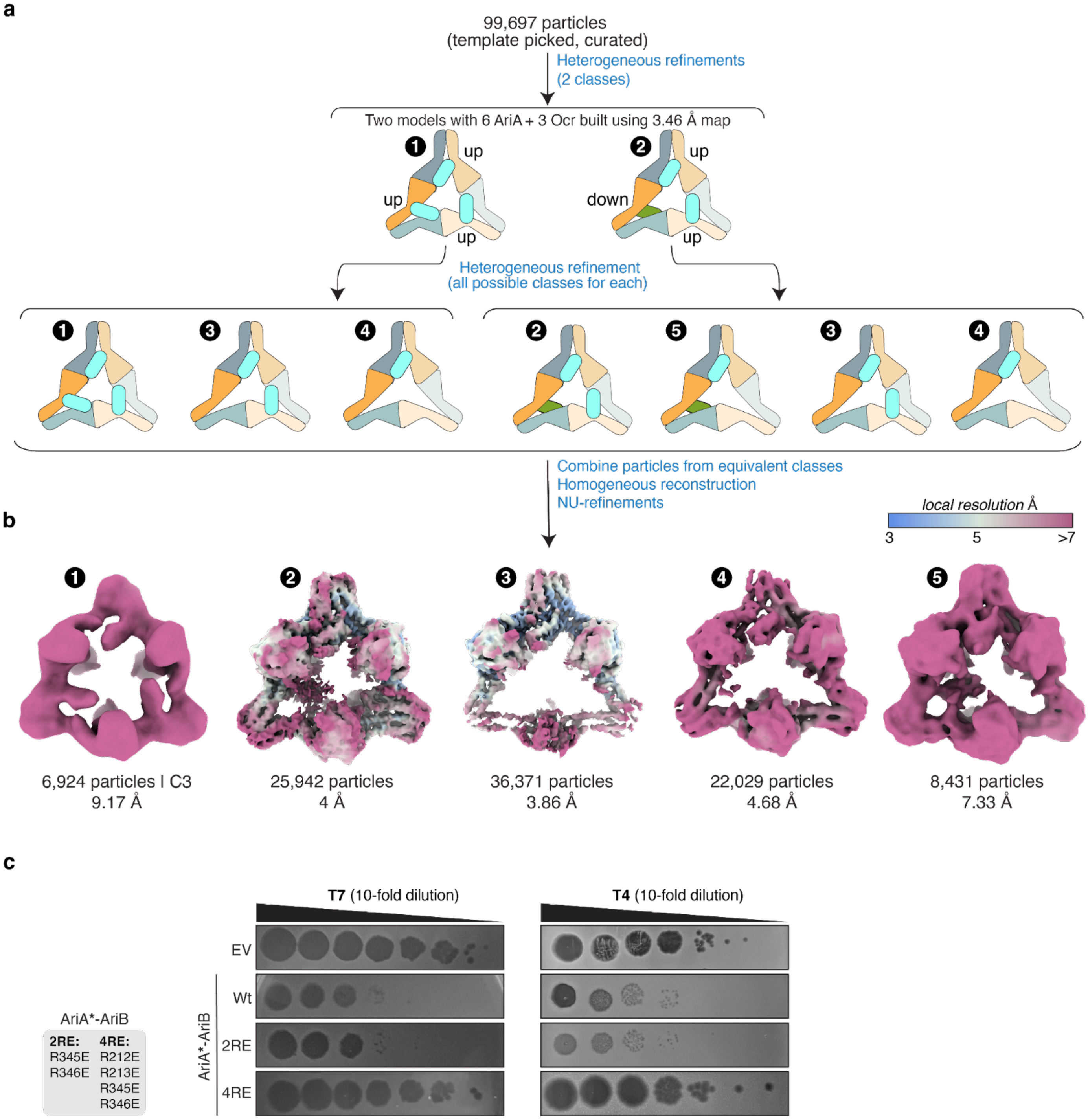


**Supplementary Figure 13 | CryoEM data processing workflow used in sorting heterogeneous AriA-Ocr complexes. (a)** Systematic sorting of compositionally and conformationally heterogeneous AriA-Ocr complexes. The cartoon schematics show the possible arrangements of AriA and Ocr subunits. Volume templates were generated using subunit structures built from the maps shown in **Supplementary Figure 12**. These volumes were imported into cryoSPARC and included in the "Heterogeneous refinement" jobs for particle class reassignment. **(b)** The final five cryoEM reconstructions for the AriA-Ocr complex, color-coded by local resolution from <3 Å (blue) to >7 Å (red). The final resolution and the number of particles corresponding to each complex form are noted at the bottom of each map. **(c)** Representative plaque-forming unit assays, demonstrating T7 and T4 phage plaques on the *E. coli* MG1655 bacterial lawn carrying either an empty vector (EV), the PARIS system (Wt; AriA-AriB as an operon), or AriA receptor pocket mutants 2RE and 4RE. 2RE and 4RE are the double and quadruple charge-reversal mutants of AriA in AriA-AriB operon (as shown in highlighted box on left). Phage 10-fold dilutions are spotted as indicated with a gradient.

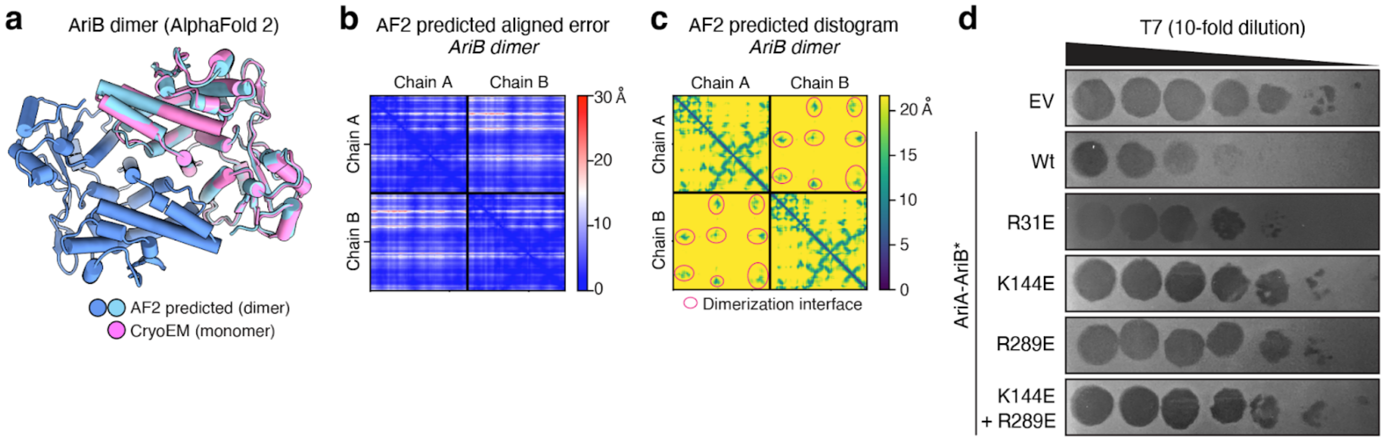


**Supplementary Figure 14 | AlphaFold structure prediction of AriB dimer. (a)** Cartoon view of the *E. coli* B185 AriB homodimer as predicted by AlphaFold 2 (blue and cyan), with one protomer overlaid with the structure of $AriB^{E90A}$ determined by cryoEM (pink). **(b)** AlphaFold 2 predicted aligned error (PAE) plot for the AriB dimer structure prediction. **(c)** AlphaFold 2 predicted distogram plot for the AriB dimer structure prediction. Red ovals indicate regions of AriB involved in the dimer interface. **(d)** A representative plaque-forming unit assay, demonstrating T7 phage plaques on the *E. coli* MG1655 bacterial lawn carrying either an empty vector (EV), the PARIS system (Wt; AriA-AriB as an operon), or AriB dimerization interface mutants: $AriB^{R31E}$ (AriA-$AriB^{R31E}$), $AriB^{K144E}$ (AriA-$AriB^{K144E}$), $AriB^{R289E}$ (AriA-$AriB^{R289E}$), $AriB^{K144E+R289E}$ (AriA-$AriB^{K144E,R289E}$). T7 phage 10-fold dilutions are spotted as indicated with a gradient.

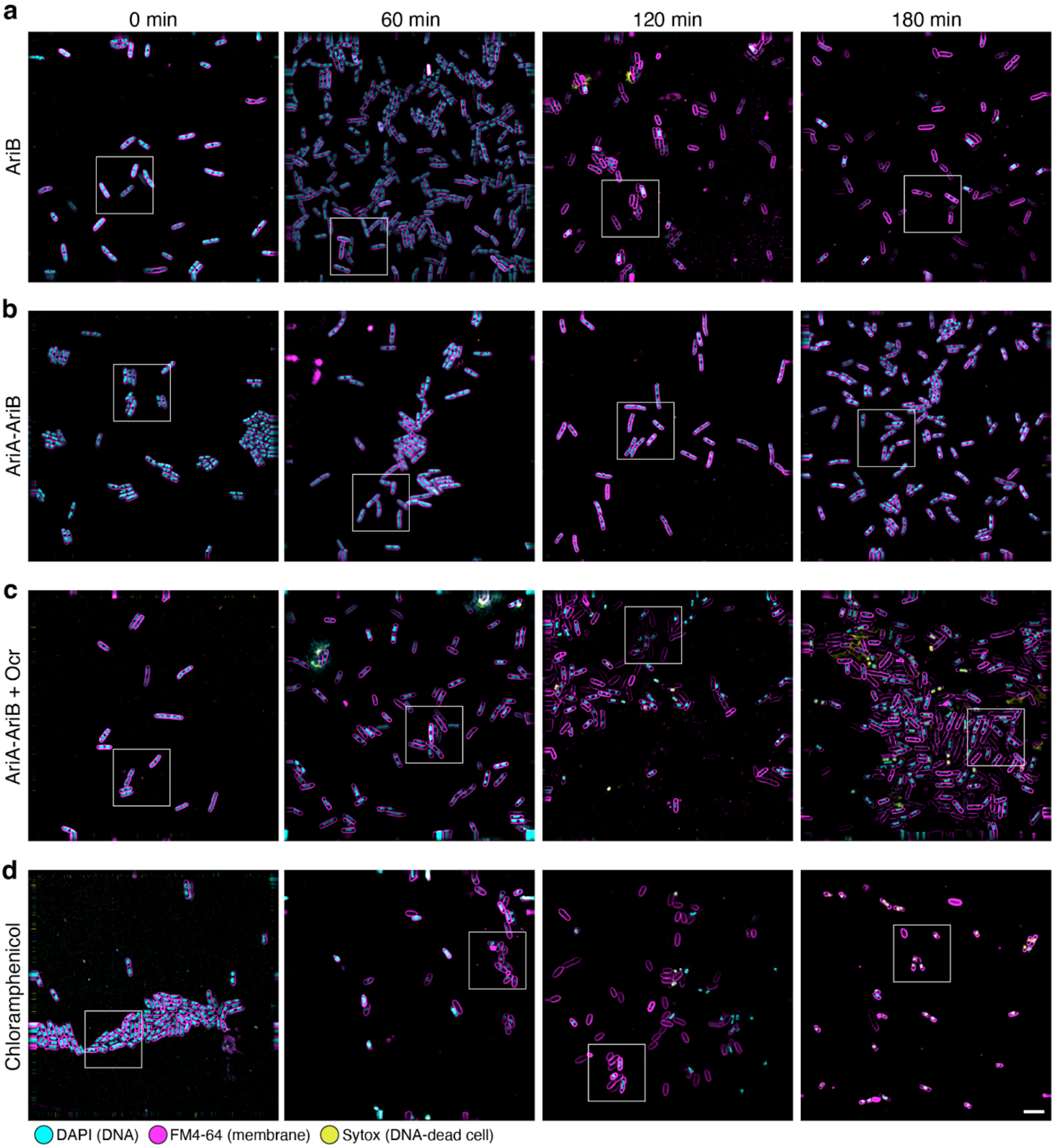


**Supplementary Figure 15 | Cytological profiling of PARIS-expressing bacterial cells. (a, b, c)** Fluorescence microscopy of *E. coli* MG1655 cells expressing AriB (panel **a**) AriA-AriB (panel **b**), or AriA-AriB + Ocr (panel **c**), at 0, 60, 120, and 180 minutes post-induction. DAPI (DNA dye) is shown in cyan, FM4-64 (membrane dye) is shown in magenta, and Sytox (DNA dye specific for dead cells) is shown in yellow. Square boxes indicate close-up views shown in **Figure 6a-c**. **(d)** Fluorescence microscopy of *E. coli* MG1655 cells treated with chloramphenicol. Colors are the same as panels **a-c**. Square boxes indicate close-up views shown in **Figure 6d**. Scale bar = 5 µm.

**Supplementary Table 1 | CryoEM data collection, refinement, and validation statistics.**

| | AriA$^{EQ}$ (EMDB-42969) (PDB 8V49) | | AriA$^{EQ}$AriB$^{E90A}$ (EMDB-42966) (PDB 8V46) | AriA$^{EQ}$AriB$^{E90A}$ (EMDB-42967) (PDB 8V47) | AriA$^{EQ}$AriB$^{E90A}$ (EMDB-42968) (PDB 8V48) | | AriA-Ocr (EMDB-42965) (PDB 8V45) | |
|---|---|---|---|---|---|---|---|---|
| **Data collection and processing** | | | | | | | | |
| Magnification | 130,000 | | 130,000 | | | | 130,000 | |
| Voltage (kV) | 300 | | 300 | | | | 300 | |
| Electron exposure (e–/Å$^2$) | 50 | | 50 | | | | 50 | |
| Defocus range (μm) | -0.5 – -2.0 | | -0.8 – -2.2 | | | | -0.8 – -2.2 | |
| Pixel size (Å) | 0.935 | | 0.935 | | | | 0.935 | |
| Symmetry imposed | C1 | | C1 | | | | C1 | |
| Initial particle images (no.) | 300,940 | | 574,500 | | | | 174,658 | |
| Final particle images (no.) | 106,506 | | 132,804 | 22,415 | 24,138 | 132,804 | 99,697 | 99,697 |
| Map resolution (Å) | 3.62 | 3.45 | 3.09 | 4.08 | 3.68 | 2.95 | 3.63 | 3.46 |
| FSC threshold | 0.143 | 0.143 | 0.143 | 0.143 | 0.143 | 0.143 | 0.143 | 0.143 |
| Map resolution range (Å) | 5.53 – 8.10 | 5.21 – 8.32 | 4.62 – 7.08 | 6.55 – 9.72 | 6.27 – 9.16 | 3.44 – 7.68 | 5.09 – 8.37 | 4.22 – 8.66 |
| **Refinement** | Global | Local | Global **Form I** | Global **Form II** | Global **Form III** | Local | Global | Local |
| Initial model used (PDB code) | *de novo* | *de novo* | *de novo* | 8V46 | 8V46 | *de novo* | 8V46, 1S7Z | *de novo* |
| Model resolution (Å) | 3.6 | 3.4 | 3.1 | 4.0 | 3.7 | 2.9 | 3.6 | 3.4 |
| FSC threshold | 0.143 | 0.143 | 0.143 | 0.143 | 0.143 | 0.143 | 0.143 | 0.143 |
| Model resolution range (Å) | | | | | | | | |
| Map sharpening *B* factor (Å$^2$) | 99.2 | 96.4 | 78.4 | 47.4 | 41.1 | 77.4 | 91.7 | 87.5 |
| Model composition | | | | | | | | |
| Non-hydrogen atoms | 12,836 | 6,431 | 15,639 | 20,588 | 25,345 | 7,579 | 19,833 | 12,076 |
| Protein residues | 1,586 | 793 | 1,937 | 2,548 | 3,141 | 936 | 2,485 | 1,492 |
| Ligands: ATP ($Mg^{2+}$) | 1 | 1 | 2 (1) | 4 | 6 | 2 (1) | 4 | 4 |
| *B* factors (Å$^2$) | | | | | | | | |
| Protein | 191.28 | 144.06 | 144.92 | 222.09 | 187.88 | 120.67 | 169.68 | 153.21 |
| Ligand | 135.59 | 124.01 | 79.43 | 176.87 | 132.68 | 119.48 | 136.86 | 151.42 |
| R.m.s. deviations | | | | | | | | |
| Bond lengths (Å) | 0.003 | 0.004 | 0.004 | 0.006 | 0.006 | 0.004 | 0.005 | 0.005 |
| Bond angles (°) | 0.795 | 0.825 | 0.588 | 0.786 | 0.733 | 0.725 | 0.751 | 0.688 |
| Validation | | | | | | | | |
| MolProbity score | 1.98 | 1.86 | 1.50 | 1.71 | 1.69 | 1.06 | 1.85 | 1.83 |
| Clashscore | 5.12 | 5.08 | 3.85 | 5.49 | 4.97 | 2.19 | 4.36 | 4.02 |
| Poor rotamers (%) | 3.08 | 2.30 | 2.58 | 2.85 | 2.49 | 0.48 | 5.55 | 4.77 |
| Ramachandran plot | | | | | | | | |
| Favored (%) | 95.19 | 95.45 | 97.94 | 97.71 | 97.33 | 97.71 | 97.70 | 97.35 |
| Allowed (%) | 4.81 | 4.55 | 2.06 | 2.29 | 2.67 | 2.29 | 2.30 | 2.65 |
| Disallowed (%) | 0 | 0 | 0 | 0 | 0 | 0 | 0 | 0 |
